# Widespread DNA off-targeting confounds studies of RNA chromatin occupancy

**DOI:** 10.1101/2025.11.11.687850

**Authors:** Micah Jonathan Goldrich, Louis Delhaye, Sarah-Lee Bekaert, Bieke Decaesteker, Filip Van Nieuwerburgh, Frank Speleman, Sven Eyckerman, Pieter Mestdagh, Igor Ulitsky

**Author notes:** co-first authors (Micah Jonathan Goldrich, Louis Delhaye).

## Abstract

The importance of long noncoding RNA (lncRNA) functions is recognized across biological systems, but their modes of action remain poorly understood. One mechanism proposed to be particularly common is gene expression regulation via recruitment to specific genomic regions. Several high-throughput sequencing methods were developed for studying the genome-wide chromatin occupancy of lncRNAs, including ChIRP-seq, CHART-seq, and RAP-seq. These methods utilize biotin-labeled probes targeting the RNA of interest to isolate and recover the chromatin associated with it. Many of the datasets obtained with these methods contain thousands of binding sites, which appears to be in contradiction with the low abundance of the interrogated lncRNAs. We studied the chromatin interactome of NESPR lncRNA in cells with varying levels of endogenous expression and then performed a meta-analysis using dozens of RNA-chromatin interaction datasets in human and mouse cells. We demonstrate that thousands of regions reported to bind lncRNAs most likely arise from the spurious recovery of DNA elements, where the ends of the recovered DNA fragments exhibit partial complementarity with the probes used for the pulldown. In addition, crucial controls were rarely used in studies profiling RNA-chromatin interactions. Therefore, most chromatin regions reported as bound by *trans*-acting RNAs in recent studies in mammalian cells appear to be technical artifacts. We provide suggestions for assessing the quality of RNA-chromatin datasets and their improvement.

## Introduction

Long non-coding RNAs are a class of non-coding RNAs longer than 200 nucleotides and molecularly indistinguishable from mRNAs. High-throughput sequencing unveiled an extensive repertoire of lncRNAs transcribed in the human genome^1^. Efforts at characterizing lncRNAs have highlighted the cellular roles and physiological relevance of some lncRNAs, but for the vast majority, these remain unclear. One mode of action proposed to be shared by many lncRNAs is the modulation of chromatin, which affects gene expression^2^. One prominent example of a lncRNA regulating chromatin is the dosage compensation of sex chromosomes by the *XIST* lncRNA, which spreads across the silenced X-chromosome and recruits chromatin modifiers^3^. Other lncRNAs were proposed to resemble *XIST*’s mode of action and to help target chromatin modification machinery to various genomic regions in *cis* or in *trans*. Therefore, discovering where lncRNAs bind to chromatin is key to understanding their function. Several methods have been developed over the past decade to reveal the genomic binding sites of lncRNAs, including ChIRP-seq^4^, CHART-seq^5^, and RAP-seq^6^. These methods rely on the crosslinking of RNA to chromatin and DNA fragmentation by sonication, followed by the pull-down with probes designed for hybridization to the lncRNA of interest, analogous to the use of antibodies to specifically recover DNA regions bound by transcription factors or histones in ChIP-seq^7^. In all these methods, sample processing includes crosslinking, DNA fragmentation, probe hybridization, probe capture, and lastly, elution from the probes. RNase is employed to release the captured DNA, thus ensuring it is RNA-bound.

Despite their conceptual similarities, the methods differ in several technical details, most prominently in probe selection and design. In ChIRP-seq, a set of 20 nt biotinylated DNA probes are commonly used to cover different target RNA regions. Methods designed for FISH probes are used to eliminate probes with significant complementarities with other genomic regions. In CHART-seq, slightly longer DNA antisense probes (~25 nt) are first experimentally tested by RNase H and filtered to retain only those that can hybridize to the lncRNA target. RAP-seq employs considerably longer antisense DNA probes (> 60 nt) covering a considerable length of the transcripts. ChIRP-seq - the first method published - has been the most popularly adopted method so far (**Tables S1** and **S2**), possibly owing to the fewer and shorter probes needed relative to RAP-seq, and since there was no recommendation to pre-select probes by a laborious experimental procedure.

These methods have been widely applied to infer lncRNA binding sites of dozens of lncRNAs (**Table S1**). In many studies, thousands of binding sites were reported for some lncRNAs in a seeming discrepancy with their low reported copies per cell, raising questions about this mode of regulation^8–10^.

Here, we first set out to study the chromatin occupancy of the NESPR lncRNA using ChIRP-seq and find that while it appears to occupy many genomic regions, these sites are recovered even when the RNA is not expressed in cells. We then set out to investigate the dozens of published RNA chromatin occupancy studies and find that a combination of insufficient experimental rigor and controls, as well as artifactual signals arising from probe hybridization to DNA, most likely at single-stranded ends of sonicated DNA molecules, result in datasets confounded by a high fraction of false positive peaks. These analyses raise substantial doubts about the prevalence of recruitment of lncRNAs to distal chromatin sites.

## Results

### ChIRP-seq suggests *NESPR* acts as a trans-acting chromatin-associated lncRNA

*NESPR* is a lncRNA expressed during differentiation of cells of the sympathetic nervous system. It has been shown to modulate PHOX2B expression by *cis-*acting genetic regulation in visceral motor neurons^11^. In addition, *NESPR* is highly expressed in neuroblastoma^12^, a pediatric malignancy of the sympathetic nervous system. To study potential trans-acting regulation by *NESPR*, we performed ChIRP-seq with a probe set designed at *NESPR*’s second exon to identify its genome-wide DNA binding sites, using *lacZ*-targeting probes as a negative control (**Fig. 1a**). Our initial experiments were conducted in IMR-32, a neuroblastoma cell line characterized by high *NESPR* expression. RT-qPCR confirmed *NESPR* enrichment with the on-target probes, but not with the *lacZ* probes (**Fig. S1a**). Peak calling with MACS2 identified 3,296 genomic binding sites. Given that IMR-32 is a *MYCN*-amplified cell line with 75–90 copies of a 3 Mb region encompassing the *MYCN* locus^13^, we omitted peaks overlapping the *MYCN* amplicon due to unreliable peak detection in this region, resulting in 2,896 binding sites (**Fig. 1a, b**). The highest-scoring peak was located at the *NESPR* locus itself, overlapping its second exon (**Fig. 1c**), which contained the region used for the design of all the capture probes. We reasoned that this signal either represented nascent *NESPR* transcripts still bound to the genomic template, local binding of *NESPR* at its own locus, or an RNA-independent enrichment of DNA due to probe complementarity with the *NESPR* locus. To explore NESPR’s potential *trans*-acting regulatory roles, we analyzed the 2,896 binding sites using both HOMER^14^ and STREME^15^ motif enrichment tools. We retrieved highly significant *de novo* motifs (**Fig. 1d**), with the most significant HOMER motif present in 791 (27.31%) peaks. Notably, we found that these *de novo* motifs aligned with the *NESPR* transcript and clustered at distinct positions (**Fig. 1e**), suggesting that *NESPR* may bind the genome through sequence complementarity, consistent with the formation of regulatory R-loops^16^. To determine whether these potential binding events regulate nearby gene expression, we stably integrated a short hairpin RNA (shRNA) targeting *NESPR* in IMR-32 cells, achieving a 50% knockdown (**Fig. S1b**). To examine whether significantly differentially expressed genes (DEGs; adj. P ≤ 0.05) were associated with *NESPR* binding sites, we assessed the ChIRP-seq coverage in a 10 kb window up- or downstream of the transcription start site (TSS) of each DEG (2,142 genes; adj. P ≤ 0.05, |log2FC| ≥ 0.5) but did not observe any clear enrichment (**Fig. S1c**). Moreover, genes that did have a nearby ChIRP-seq peak, such as *GCH1* and *BICC1* (**Fig. 1b, Fig. S1d**), were not differentially expressed. For DEGs with a moderate expression change (2,142 genes; |log2FC| ≥ 0.5), 2.3% (122 genes), 2.3% of DEGs were located within 10 kb of a *NESPR* peak. For DEGs with a strong expression change (224 genes, |log2FC| ≥ 1), only 3 DEGs were found within 10 kb of a *NESPR* peak. Permutation testing revealed that the observed proximity was not significant for either moderate or strong DEGs (P = 0.08 and P = 0.28, respectively), indicating that the proximity of these genes to *NESPR* peaks was not greater than expected by chance (**Fig. S1e**). Similar results were observed when considering a 50 kb window around the TSS, with 7.7% of moderate DEGs (165 genes; P = 0.08) and 8.6% of strong DEGs (19 genes; P = 0.41) located near *NESPR* peaks.

**Figure 1.**
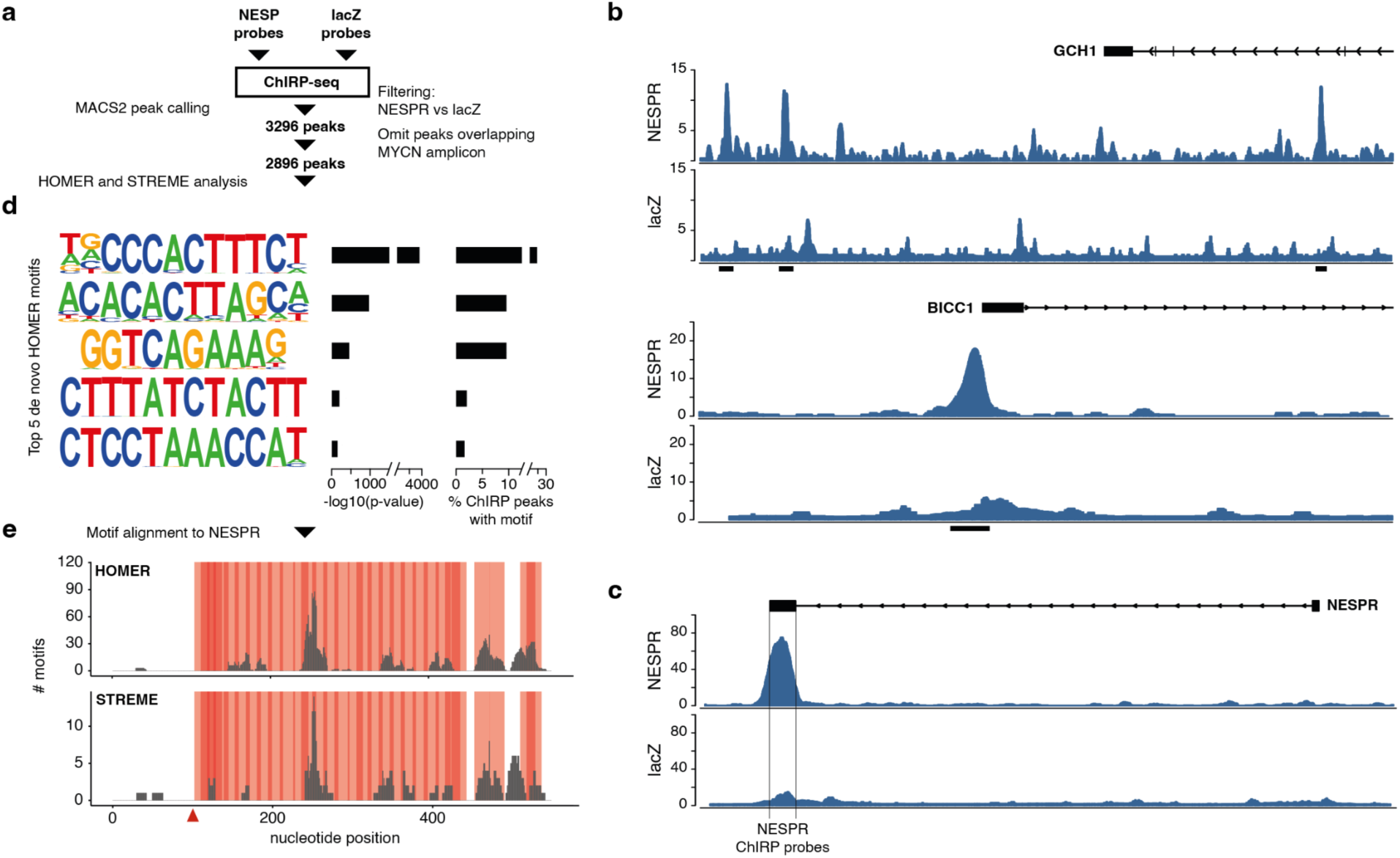
ChIRP-seq suggests a *trans*-acting function for *NESPR*. a, Overview of the peak discovery from *NESPR* ChIRP-seq in IMR-32 cells. b, ChIRP-seq peaks at *GCH1* and *BICC1* loci. c, ChIRP-seq peaks at the *NESPR* locus itself. d, Top 5 *de novo* HOMER motifs enriched in the unique NESPR peaks. e, Alignment of all *de novo* HOMER and STREME motifs to the *NESPR* transcript per nucleotide. Red bars highlight the tiling probe positions on the *NESPR* transcript. Red arrow shows the splice junction.

### RNA-independent genomic DNA enrichment confounds *NESPR* ChIRP-seq data

Given the lack of statistical evidence for the functionality of *NESPR* DNA-binding on gene expression, we considered an alternative explanation. We hypothesized that the *NESPR*-specific probes might also directly capture genomic DNA, regardless of *NESPR* RNA enrichment. To test this hypothesis, we repeated the ChIRP-seq experiments using additional controls (Fig. 2a), including an RNase-treated lysate, as well as a lysate from SHEP cells, which do not express *NESPR* (Fig. S2a). Enrichment in IMR-32 was confirmed by RT-qPCR, and as expected no enrichment was observed in RNase-treated IMR-32 and SHEP lysates (Fig. S2b). Following stringent filtering against input, RNase-treated IMR-32, and SHEP cell backgrounds, only 3 genomic binding sites were retained, in contrast to the 2,896 peaks identified in the original analysis using only *lacZ* probes as a negative control. None of the retained peaks overlapped the *MYCN* amplicon. As before, the *NESPR* locus itself was enriched in IMR-32 cells, however similar enrichment was also observed in both the RNase-treated IMR-32 and *NESPR*-negative SHEP controls (Fig. 2b), suggesting that this signal is independent of *NESPR* RNA and instead reflects direct probe hybridization to the genome. Moreover, most peaks identified in our original dataset (IMR-32) were also detected independently across all three new ChIRP-seq experiments (IMR-32, IMR-32 RNase and SHEP) (Fig. 2c, d), reinforcing the conclusion that these regions are reproducibly recovered independently of *NESPR* RNA.

**Figure 2.**
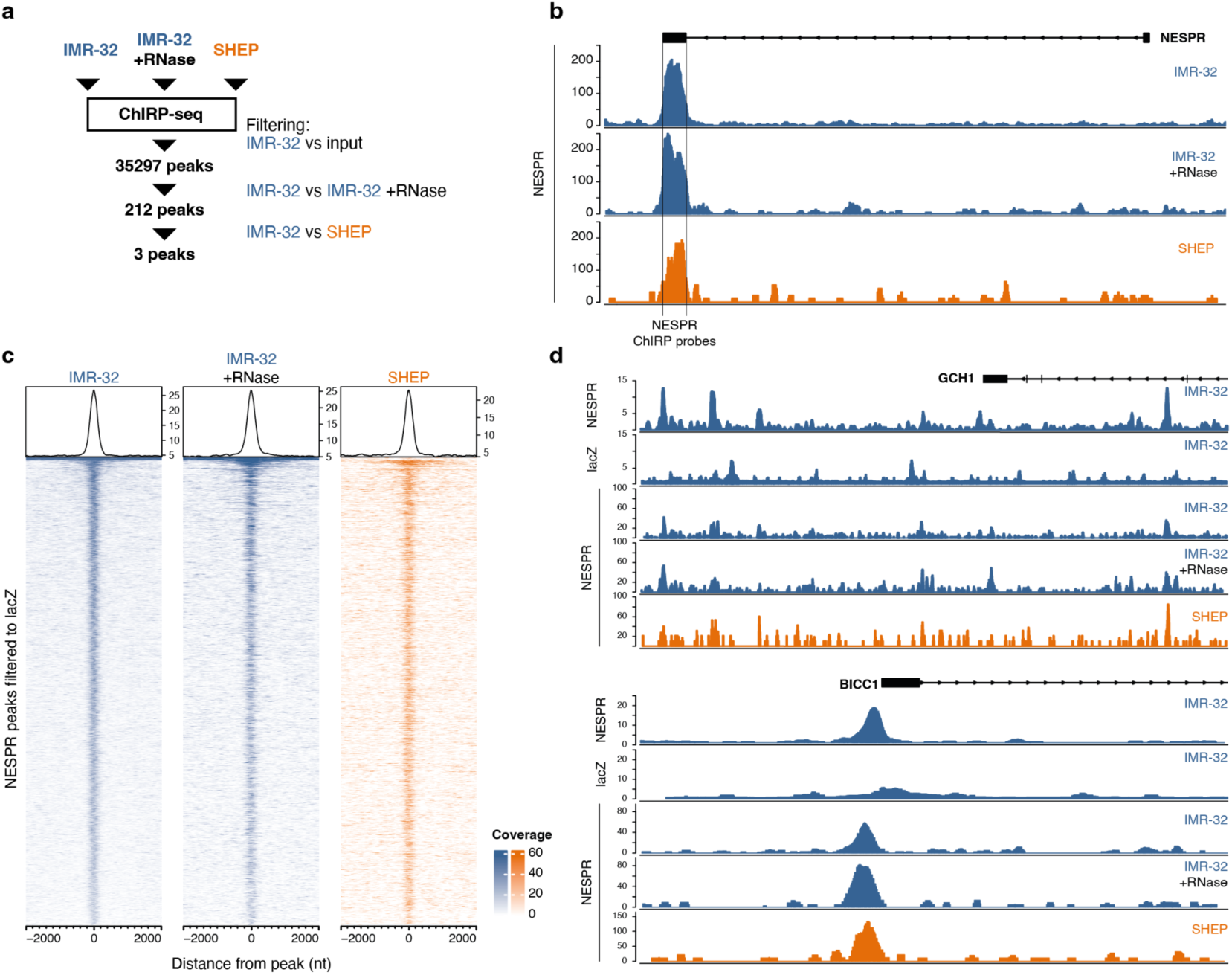
Stringent experimental controls demonstrate genomic DNA enrichment independent of *NESPR* RNA enrichment. a, Revised filtering strategy for *NESPR* ChIRP-seq. b, ChIRP-seq peaks at the *NESPR* locus. c, Coverage of the revised ChIRP-seq experiments at the previously retrieved 2,896 peaks. d, ChIRP-seq peaks at the *GCH1* and *BICC1* loci. Dark blue, cells with *NESPR* expression; orange, cells without *NESPR* expression.

### A compendium of RNA-chromatin interaction studies

To evaluate more broadly RNA-chromatin interactions sequencing datasets we gathered data from 22 human and 38 mouse studies, using ChIRP-seq, CHART-seq, or RAP-seq **(Table S1**). All human studies and a majority of mouse studies were performed using the ChIRP-seq protocol. The remaining mouse studies used the CHART-seq, RAP-seq and CHIRT-seq (a combination of ChIRP-seq and CHART-seq), with Xist being the gene of interest (GOI) in 8 of the 13 non-ChIRP studies (and the only GOI in all RAP-seq experiments). We established and employed a bioinformatic pipeline based on Bowtie2^17^ for read alignment and MACS2^18^ for peak calling, a combination of tools that was used in 16/18 human studies that reported peak calling, to process all datasets uniformly. We also obtained the peak sets reported in the original studies, which were often derived from a computational integration of the ‘even’ and ‘odd’ peak sets (described below).

A comparison of several experimental outlines revealed clear differences among the studies (**Fig. 3a** and **Table S1**). As in other pull-down–based methods, the original ChIRP-seq publication suggested an input sample to be used as background to control for sample bias which may arise from such things as genome aberrations, repeat expansions, or experimental condition artifacts. Of note, 6 and 5 human and mouse studies, respectively, had no such input samples sequenced and/or published. Biological repeats of the same sample and probe set are the only means to evaluate internal reproducibility, yet biological replicates were performed in only 9/22 human studies, and in 20/38 mouse ones. Thus, many studies concluded on chromatin occupancy patterns of an RNA based on a single biological replicate. Also, among the human and mouse studies with biological replicates, only seven studies performed three repeats per experiment.

**Figure 3.**
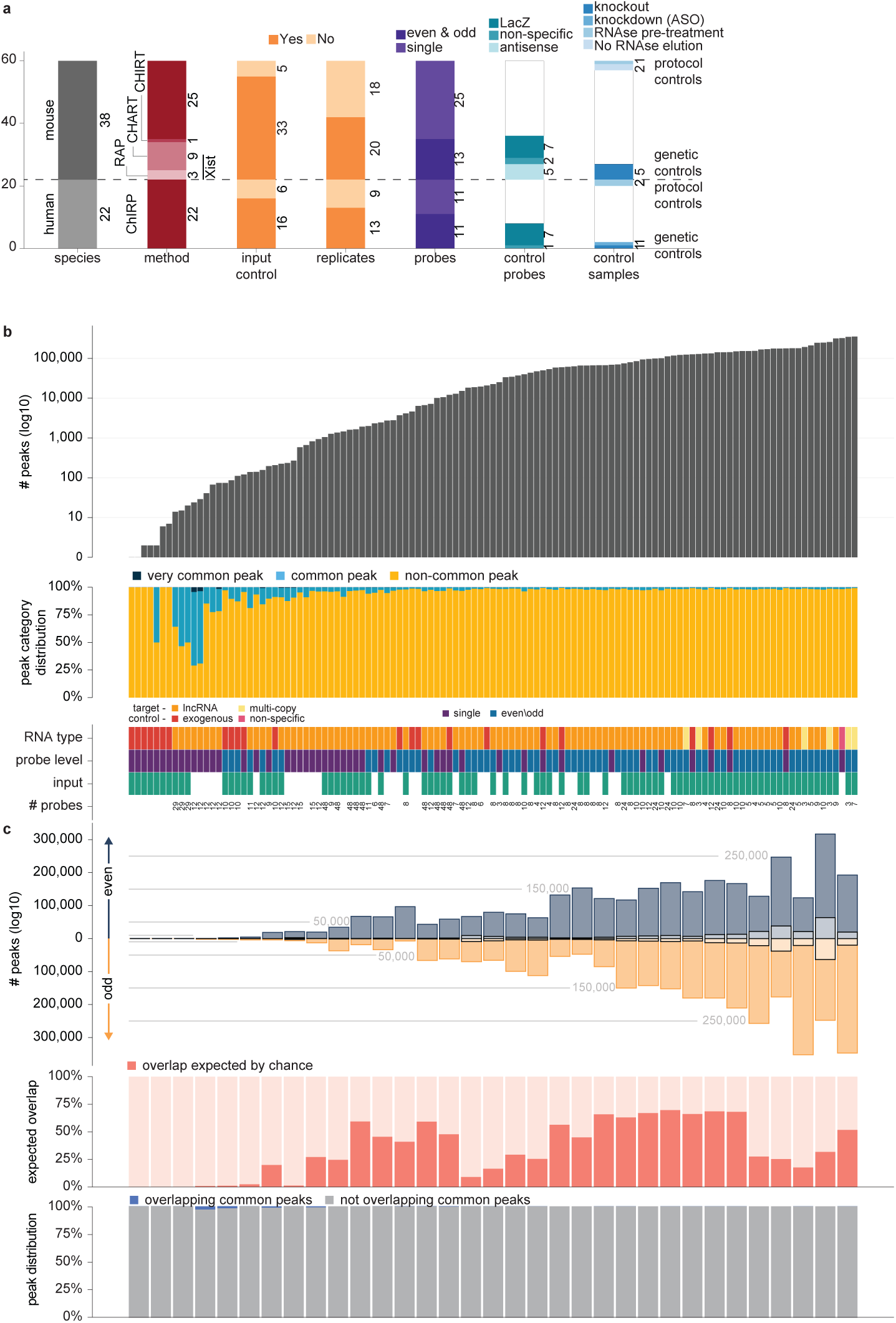
Peak distributions and properties across human samples, including matched replicates. **a,** Characteristics of RNA-chromatin occupancy studies are characterized by (order from left to right): species; experimental protocol; presence of input samples in study; replicates performed; probe types; use of control probe sets; use of control samples. **b,** Top: Total number of peaks detected per sample, sorted in ascending order. Middle: Proportion of peaks assigned to three abundance categories: *non-common*, *common* (detected in 6≥ studies), and *very common* (detected in ≥10 studies). Bottom: Summary of probe and experiment-related variables for each sample, including the RNA target class, number of probes (when available), probe pool type, and presence or absence of a complementary input sample. **c,** Analysis of replicates using matched “even” and “odd” probe pools. Top: Barplot showing the number of peaks in the “even” (top) and “odd” (bottom) replicates for each matched pair. Overlapping peaks between replicates are shown as a transparent overlay. Middle: Observed overlap between replicates expressed as a percentage of the overlap expected by chance, based on random peak placement. Absent overlapping peaks are striped red and white lines. Bottom: Abundance classification of the overlapping peaks (non-common, common, very common) based on their recurrence across the entire dataset. Absent overlapping peaks are striped red and white lines.

Binding of target probes to off-target DNA due to partial probe complementarity is a potential pitfall in ChIRP-seq data that was already recognized in the original publication of the method^19^. To control for such non-specific probe–DNA binding, the authors proposed the splitting of the target probes into two probe sets termed ‘even’ and ‘odd’, such that only the intersection of peaks obtained from both sets of probes should correspond to true binding of the lncRNA to a DNA region. Nonetheless, a single “on target” probe set was used for 12 of the 25 GOIs in human studies and for 22 of the 41 genes in mouse studies (of these 22, 11 are RAP-seq, CHART-seq and CHIRT-seq studies). We note that a single set was also used for *NESPR*, due to its short length. In addition, several studies utilized control probe sets of a different nature, including probes targeting an exogenous gene, non-specific controls (random or targeting a different gene), and control probes antisense to the true probe set (which therefore cannot hybridize to the RNA target, but can potentially still bind similar DNA regions, see below). A total of eight studies in human and 14 studies in mouse employed some control probe sets, with 14/22 using only *lacZ* probes (that are not expected to recover any RNA).

Antisense control probes, which as we will discuss below are potentially particularly useful, were only used in CHART-seq studies. Overall, control samples were sporadically used in the analysis in their respective publications. For example, half of the publications that used *lacZ* probes did not indicate at all how they were used in the analysis or what the *lacZ*-probe–recovered peaks mean for their data interpretation.

Finally, very few studies used controls that would reveal the RNA dependence of the recovered DNA sequences, specifically genetic manipulations or protocol changes (**Fig. 3a**). In human studies, one study used GOI knockout cells and another study performed GOI knockdown, and five mouse studies analyzed GOI knockout cells. In terms of protocol controls, sample treatment with RNase prior to probe incubation – thus removing RNA from the sample – was performed in two studies, and a control with no RNase elution was performed in a single human study and two mouse ones.

Overall, RNA-chromatin interaction assays have been performed with greatly varying conditions and controls, and in most cases ChIRP-seq was performed using fewer probes and controls relative to the original publication.

### Large variation in the number of detected genomic target regions among studies and samples

Ultimately, the goal of RNA-chromatin interaction mapping is to identify sites that are bound by the RNA of the GOI, with the objective of deriving biological insight such as the function and mechanism of action. Therefore, we first focused on comparing these studies through the prism of the peaks identified from each dataset. We found substantial variation in peak number, with some samples having very few peaks and others having hundreds of thousands of peaks (**Fig. 3b** and **Fig. S3a** top panel). The mean number of peaks per sample in human and mouse studies is 65,700 and 30,764, respectively. We note that in the vast majority of cases the targeted RNAs are expected to be expressed at few copies per cell, at a seeming contradiction to thousands of binding sites. Notably, if the lncRNA binds its target sites in only a small subset (<1%) of the millions of cells profiled in each experiment, we wouldn’t expect this binding to result in a peak in the corresponding dataset, as such sites are expected to be indistinguishable from background.

We first examined common peaks that appeared regardless of cell-type and probe target, and may therefore represent non-specific and problematic regions. We removed reads overlapping the ENCODE blacklist, containing anomalous regions with high signal in sonicated DNA independent of cell line or experiment^20^. We merged peaks from the different samples, excluding those involving exogenous targets (e.g., luciferase RNA), multiple studies studying the same GOI (e.g., *Xist*) and studies where the GOI corresponded to a repetitive-element (e.g., LINE RNA), to avoid biasing the merged peak set. For each merged peak region, we overlapped peaks from individual samples and counted in how many distinct studies we had identified a peak within that region. Merged peaks supported by at least six studies were classified as “common peaks,” while those present in ten or more studies were designated as “very common peaks” (**Fig. S3b**).

In human datasets, 6,430 of the 2,080,068 merged peaks (0.3%) were common peaks, and only a single peak met the threshold for a very common peak. In mouse datasets, 10,970 of 1,711,140 merged peaks (0.6%) were common peaks, and 39 were very common peaks. These results indicate that, in some contrast to results from ChIP-seq and Cut&Run studies^21–23^, common and very common peaks constitute only a small fraction of all peaks observed in the profiled studies. In most studies, a minor subset of the total peak corpus was common, with notable exception of the studies with <100 peaks, where common peaks sometimes had a dominant presence, in particular in mouse (**Fig. 3b and Fig. S3a** middle panels).

### Peaks recovered by even and odd probe sets are often highly discordant

We next considered the “even” and “odd” samples recovered using the alternating sets of probes, proposed as a corrective strategy for potential direct hybridization of the biotinylated probes to DNA^24^. We first considered separately the peaks identified using each probe set, which we refer to as ‘even peaks’ and ‘odd peaks’. We will refer to each such pair as a ‘matched’ sample. Across most matched samples, the numbers of even and odd peaks can often vary by up to 3-fold and rarely exceed an order of magnitude (typically only when the total number of peaks was very low, e.g., 22 and 2; **Fig. 3c** and **Fig. S3c** top panel). More importantly, the overlap of each matched sample typically represents only a small fraction of either the “even” peaks or the “odd” peaks, indicating many peaks in each sample are unlikely to be true binding sites of the GOI. Notably, although the overlap represents a small fraction of the peaks, it is statistically significant, at least when evaluated against a simplistic random model that shuffles peak positions in each chromosome (**Fig. 3c and Fig. S3b middle panel**).

In matched samples, common and very common peaks were less frequently observed in human studies. In contrast, mouse studies exhibited a greater number of such peaks, particularly in cases where the matched sets contained fewer total peaks (**Fig. 3c** and **Fig. S2b** bottom panel).

The low rate of overlap between even and odd peaks, raises questions as to the source of the large number of peaks obtained for most lncRNAs. We hypothesized that partial probe hybridization to complementary sequences could explain the recovery of these regions and this occurs independent of the RNA, yielding false-positive binding regions.

### Peaks commonly contain short matches to probe sequences

Since GOI-targeting probes are designed using computational tools developed for FISH experiments, and those generally exclude sequences with perfect or near-perfect matches elsewhere in the genome, we hypothesized that DNA binding may stem from shorter, partial sequence matches. To investigate this possibility, probe sequences were decomposed into *k*-mers of lengths ranging from 7 to 20 nt (**Fig. 4a**; 20 nt is the typical length for ChIRP-seq probes). For each probe-derived *k*-mer set: “even,” “odd,” (for studies with split even/odd probe sets) or “single”, we assessed their occurrences across the peak sequences of all relevant samples, recording both the number and positions of matches per probe set. To evaluate occurrences expected by chance, we generated 10 dinucleotide-shuffled versions of each probe set and analyzed them in the same manner (**Fig. 4a**). This analysis was performed across the raw peak sets of individual samples (“even,” “odd,” and “single”), as well as when considering strictly overlapping peaks from even and odd samples, either merging the intersecting even/odd peaks (“union”) or taking just their intersection region (“intersect”). When available, published consensus peak sets were also included in the analysis (**Table S3**).

**Figure 4.**
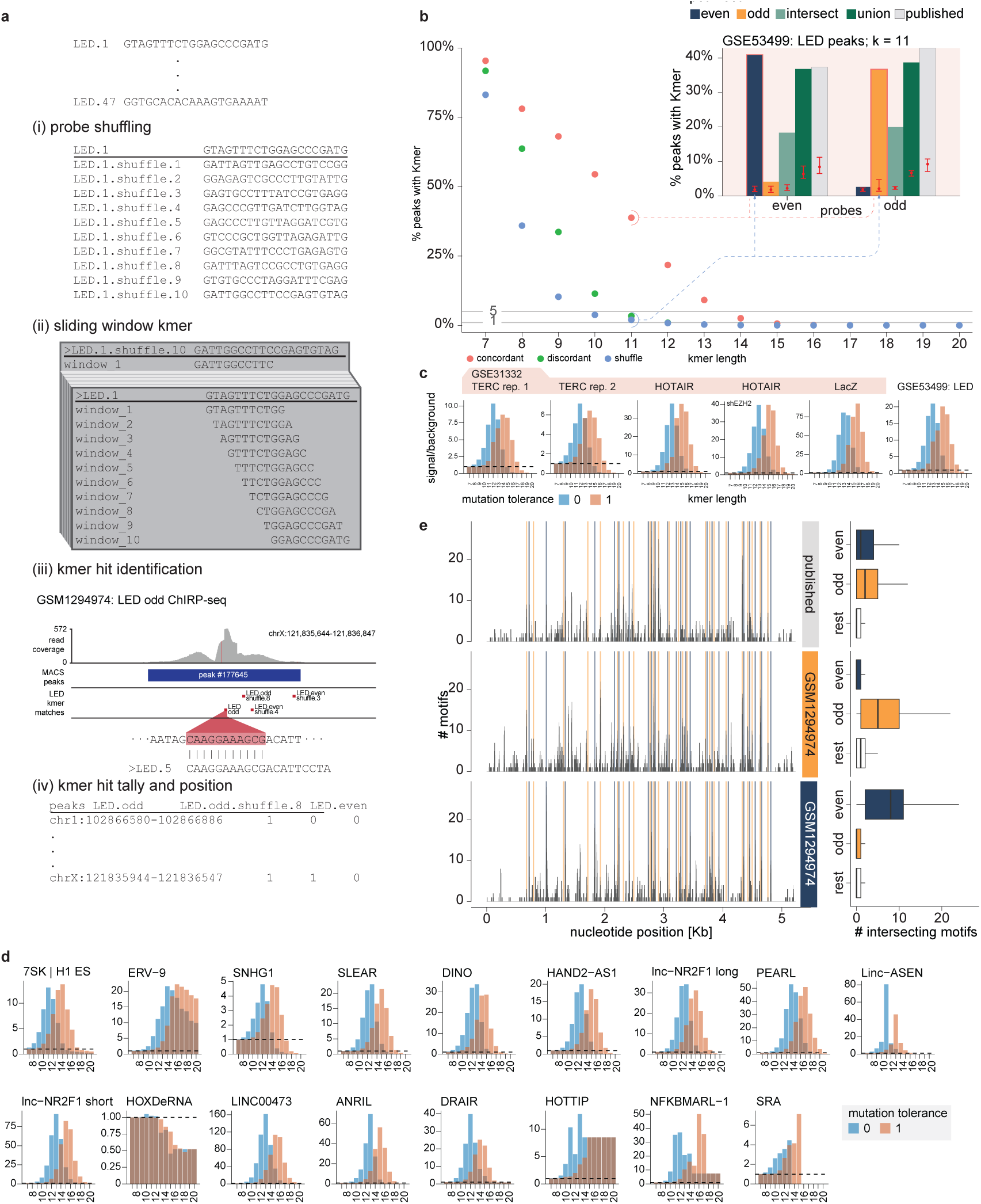
Peaks recovered in chromatin-RNA interaction studies often share kmers with the probes. **a,** Schematic outline of the procedure used to detect probe-derived k-mers: (i) shuffled probe sequences are generated as a background control; (ii) a sliding window is used to generate all possible k-mers of a given length from each probe set; (iii) k-mer matches are searched across all peak regions; (iv) the positions of matching k-mers within the peaks are tallied and recorded. **b,** Inset: Positional distribution of a representative 11-mer from the LED probes across all sample peaks. Red whiskers indicate the minimum, maximum, and dot indicates the mean levels of matching from shuffled control probes. Main: Enrichment levels of probe-derived k-mers across k-mer lengths from 7 to 20, categorized as concordant (probes matching their intended target), discordant (probes mismatched to the target), and shuffle (randomized controls). **c,** signal-to-background ratios for LED and the original ChIRP-seq study samples. **d,** Signal-over-background values are plotted for k-mers of length 7 to 20 across grouped human samples. Sample groups are ordered by the average number of peaks per group, arranged from left to right and top to bottom. Groupings were defined based on identical experimental conditions, probe target, and biological replicate number. When applicable, both “even” and “odd” probe sets were included; otherwise, single-probe sets (“single”) were used. **e,** Left: Distribution of motif coverage along the LED lncRNA transcript. Right: motif coverage across bases by probe group.

When examining the fraction of peaks with at least one *k*-mer match across the raw samples in studies with “even” and “odd” probes, a pattern emerges. In concordant combinations of probe sets and peaks (e.g., “even” peaks and “even” probes), short *k*-mers show similar match rates between real and shuffled probe sets—consistent with the high background probability of finding matches for short sequences. However, as *k*-mer length increases, a significantly larger fraction of peaks contains matches to *k*-mers found in the true probe set compared to their shuffled counterparts. This enrichment becomes evident at intermediate *k*-mer lengths (e.g., *k*=11 for the *LED* lncRNA study^25^; **Fig. 4b**, upper-right inset), and persists for longer *k*-mers, though the absolute number of matches declines as expected, due to the lower chances of finding longer exact matches in the genome.

Notably, discordant combinations (“even” peaks with “odd” probes or vice versa) do not show such enrichment for their respective *k*-mers and instead more closely match the background signal of the shuffled controls. This suggests that the observed enrichment most likely reflects sequence-specific hybridization of the individual probe sets to DNA molecules, similarly as we demonstrated for *NESPR* experimentally. In line with previous probe design constraints that avoid perfect or near-perfect matches outside the gene of interest (GOI), the remaining matches for very long k-mers often map to the genomic locus of the GOI itself (**Fig. S4A**).

Furthermore, when assessing probe enrichment in the “union,” “intersect,” and “published” peak sets, *k*-mers found in both “even” and “odd” probe sets are enriched relative to their shuffled counterparts (**Fig. 4b**). While one might hope that intersecting two independently derived peak sets would help isolate true binding events by filtering out probe-specific artifacts, this does not appear to be the case. Enrichment for both probe sets persists also in the intersected regions. This underscores that even widely used strategies for defining consensus peaks, ranging from simple set operations like intersection and union to the diverse methodologies employed in published studies, do not effectively eliminate likely RNA-independent enrichment of DNA fragments, likely because the large number of DNA fragments recovered by each probe set, inevitably results also in a substantial number of peaks that happen to contain complementary region to probes from both sets.

To quantify the extent of probe-driven spurious peaks, we calculated the ratio between the mean *k*-mer match rate in concordant combinations (i.e., “even” probes in “even” peaks, “odd” probes in “odd” peaks, or “single” probe sets in studies that did not use split probe sets) – representing the signal – and the mean match rate in their respective shuffled controls, representing the background. Plotting this ratio across all samples revealed a consistent enrichment for *k*-mers longer than 10, indicating that partial hybridization of probe sequences to genomic DNA fragments is very widespread. This pattern is evident for the *LED* lncRNA as well as for other targets from the original ChIRP-seq study, as illustrated in **Fig. 4c**. A sample of signal/background ratios for human studies is shown in **Fig. 4d** and detailed panels for all studies are in **Fig. S4B** and **S5**.

When examining where within the peaks the probe matching signal appears, we observed a consistent central enrichment across all sample types. This density becomes increasingly pronounced for intermediate to longer *k*-mers, further indicating that these sequences are likely driving peak formation through direct probe-to-DNA interactions (**Fig. S6**).

### Probe off-targeting leads to the enrichment of specific sequence motifs in peaks

The recruitment of lncRNAs to genomic sites has often been suggested to be sequence-specific and driven by sequence motifs, some of which were reported to be enriched in ChIRP-seq peaks in various studies^27–29^. For example, several long motifs derived from the *ANRIL* binding sites were shown to match sequences in the *ANRIL* transcript^30^. Such correspondence between lncRNA binding site motifs and the GOI sequence can imply a sequence-dependent binding mediated by Watson-Crick base-pairing. Notably, a recent meta-analysis revealed binding motifs corresponding to the target transcript sequences across multiple samples and suggested that RNA:DNA structures are a significant and widespread theme of lncRNA binding to chromatin^31^.

Based on our motif enrichment analysis for *NESPR*, we reasoned that an alternative hypothesis for the presence of enriched short motifs within binding peaks is by partial and RNA-independent probe hybridization to DNA. Under this model, motifs intersecting with the target sequence should be preferentially found near the positions of the regions targeted by the probes relative to other regions of the transcript. To assess this hypothesis, we performed uniform analysis of all the peak sets with the *de novo* motif discovery tools STREME^32^ and HOMER^33^, and aligned the motifs across the GOI sequence using FIMO^34^. Concordantly with the expectation of a model in which the motifs are driven by DNA hybridization with the probes and not with the lncRNA, across many lncRNAs and samples a clear enrichment of motif instances is found at the locations of the probe relative to other regions of the transcript, such as for the lncRNA *LED* (**Fig. 4e**). Motif coverage across the GOI aligns closely with the regions targeted by the probes, with a notable enrichment of motifs at the specific nucleotide positions corresponding to the probe sequences. This pattern is also evident for the *DINO* lncRNA, including in samples pre-treated with RNase, further supporting direct probe-to-DNA binding as the likely source of the signal (**Fig. S7**). Additionally, we observe that some regions covered by probes lack any intersecting motifs, suggesting that some probes might have reduced tendency to enrich complementary DNA. This may also reflect the non-uniform sequence composition of the genome, where certain k-mers are more prevalent and therefore more likely to seed motifs that appear enriched in peak sets.

Together, these analyses reveal that many peaks and identified motifs most likely arise from direct probe-to-DNA hybridization rather than genuine RNA-chromatin associations. This technical artifact inflates the apparent genomic occupancy of the RNA of the GOI and can lead to misinterpretation of motif enrichment as biological insight, when it instead reflects probe sequence bias.

### Longest *k*-mer profiles reveal composition of intersection peak sets

Ideally, strategies such as intersecting peaks across independent probe sets should enrich biologically meaningful signals while filtering out technical artifacts. Thus, while the enrichment of *k*-mers in peaks obtained using either even or odd probes may simply reflect probe binding and is not inherently problematic, its persistence in the intersection peak sets considered to correspond to true binding sites is more concerning. The continued presence of probe-matching *k*-mers in these refined sets as mentioned above suggests that the false positives are not being adequately removed. To pinpoint the potentially most influential probe-matching sequence within each peak, we introduce the concept of the “longest probe-matching *k*-mer”, defined as the longest *k*-mer found within the peak that is identical to a probe sequence (**Fig. 5a**). These longest *k*-mers show a central enrichment across peaks (**Fig. S6a**), reinforcing their likely role as drivers of peak formation.

**Figure 5.**
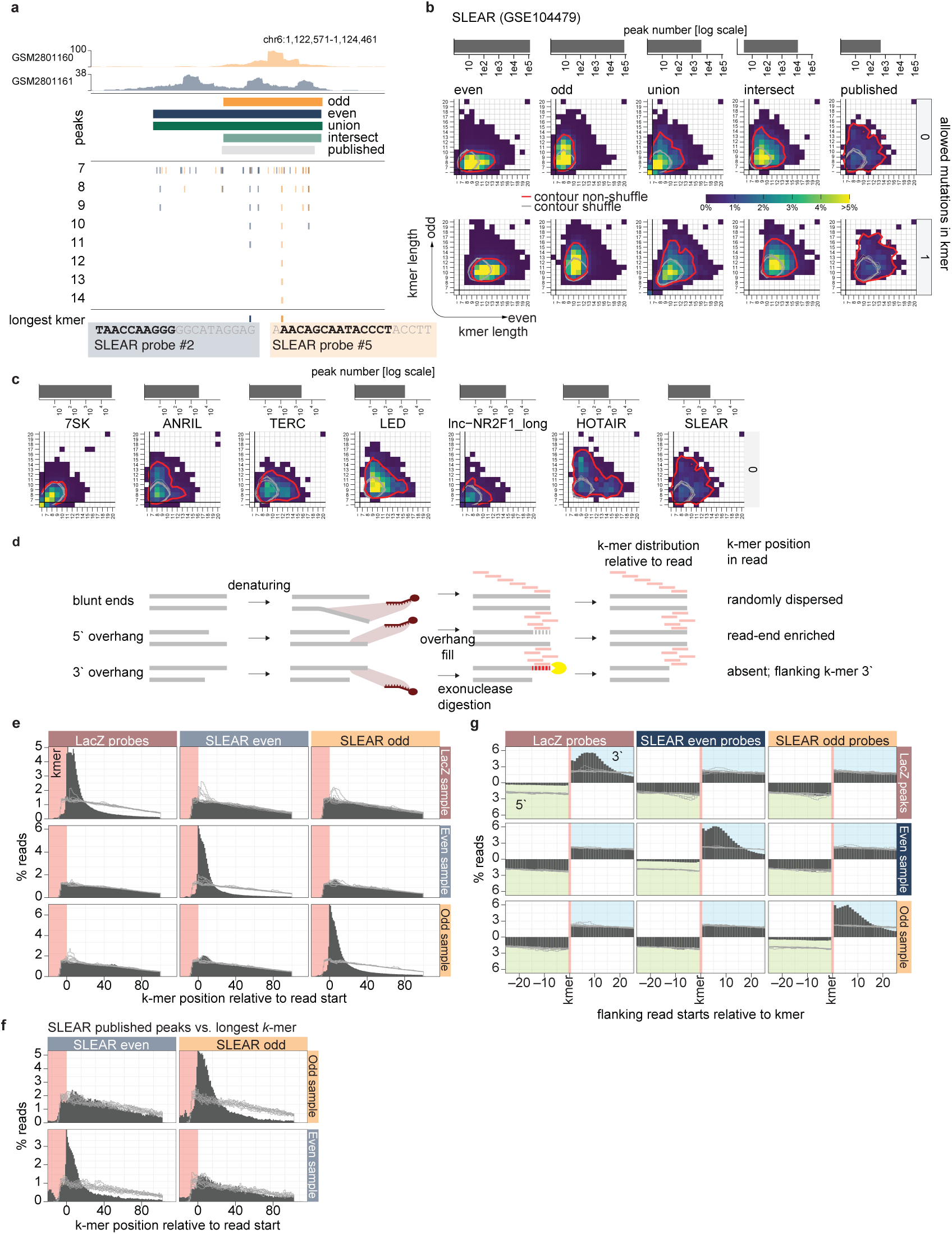
Longest k-mers and association of k-mers with read ends. **a,** Schematic outlining the identification of the longest k-mer and associated reads. **b-c,** Heatmap of even–odd k-mer pairings across peak sets. The top panel indicates the number of peaks per set. Contours delineate the top 90% density range for observed (red) and shuffled (gray) k-mer combinations. Shown are heatmaps for different peak sets for one SLEAR dataset (b) and published peaks from different studies (c). **d,** Illustration of different fragment end types and their expected impact on the positioning of k-mers within reads. **e-f,** Distribution of read start positions relative to the start of the longest k-mer across raw samples. K-mers that intersect but do not span the entire match are shaded in pink. Gray lines correspond to shuffled probes. Shown are analysis for the SLEAR study using our processing of the data (e) using published peaks (f). **g,** Aggregate signal of read start positions flanking the longest k-mer. Shaded regions highlight enrichment on the 3′ side (blue) and 5′ side (green) relative to the k-mer (pink). Gray lines correspond to shuffled probes.

For each peak set, we constructed a two-dimensional matrix, categorizing it by the longest probe-matching *k*-mer found in its “even” and “odd” probes. In the “even” and “odd” samples, enrichment for probes-matching sequences is found on the axis corresponding to the probe set, reaching up to *k*-mer length of ~13 nt, with little contribution from the opposite set (**Fig. 5b**; red contours). In contrast, for shuffled probes, most peaks fall within the short *k*-mer range with no axial preference (**Fig. 5b**; gray contours). This pattern is in line with what is seen when examining *k*-mer representation alone (**Fig. 4b**).

When examining unions or intersections of even and odd peaks, the distribution of *k*-mer combinations broadens with a strong signal peak at short *k*-mer combinations that extends outward in multiple directions. This diffuse pattern indicates that simple set operations are likely insufficient to flag peaks that result from an RNA-independent direct probe hybridization; instead, these peaks likely result from partial hybridization with both even and odd probes through spatially proximal but independent *k*-mers.

When examining peaks reported in the individual studies (**Fig. 5c** and **Fig. S8**), which used different methods to integrate data from libraries recovered using even and odd probes, the peak sets often exhibit even broader and more study-specific k-mer distributions, highlighting how different consensus-calling pipelines contain varying fractions of likely probe-driven peaks.

### Probe off-targeting preferentially occurs at the ends of recovered DNA fragments

Having established common off-targeting of DNA regions by the probes used in RNA-chromatin interaction profiling, we were interested in the mechanism underpinning this phenomenon. To this point, our analyses have centered on identifying systematic patterns of probe-derived bias at the peak level. We were next interested in understanding the molecular basis for the formation of peaks carrying probe-matching sequences. We considered two possible scenarios for how probes hybridize to DNA and drive peak formation (**Fig. 5d**). In the first scenario, probes hybridize to DNA fragment ends—either staggered and inherently exposed, or blunt ends that transiently denature—providing sufficient exposure of single-stranded DNA for probe binding. Importantly, the process of DNA sonication can indeed produce staggered DNA ends^35,36^. In the second scenario, conditions prior to or during hybridization result in widespread DNA denaturation and accessibility, allowing probe binding anywhere along the fragment. These mechanisms make distinct predictions about the location of probe-derived *k*-mers within sequencing reads: a bias toward read starts in the first case, versus uniform distribution in the second (**Fig. 5d**).

When mapping the position of the longest *k*-mers within their intersecting sequencing reads, we observe a strong bias toward the read starts for true probe sets (**Fig. 5e**). This pattern disappears in control comparisons using shuffled probes or discordant probe-peak combinations, which show a rather uniform distribution (**Fig. 5e**). These observations indicate that probes preferentially bind near the start of sequencing reads, implicating DNA ends as the key access points for probe hybridization.

Notably, the same positional bias is also observed in longest *k*-mer–containing reads derived from published peak sets (**Fig. 5f**). This pattern extends across a wide range of peak numbers, including several CHART-seq datasets—suggesting that the same mechanism for probe hybridization is applicable across methods.

To validate this interpretation, we further leveraged a technical nuance of sequencing library preparation. DNA fragments generated by sonication often have staggered ends that must be converted to blunt ends prior to sequencing. While 5′ overhangs can be filled in and retained in sequencing reads, 3′ overhangs are digested and thus removed. This asymmetry provides a testable prediction: if probes bind to 3′ overhangs, the corresponding *k*-mer will not appear in the read itself, but the read will begin just downstream of it (**Fig. 5g**). Indeed, we observe an enrichment of read starts immediately downstream (3′) of the longest *k*-mer locations—consistent with digestion of 3′ overhangs—while no such enrichment is observed upstream (5′) or in control datasets. This asymmetry further strengthens the evidence that probe binding preferentially occurs at fragment ends and those staggered ends, in particular, are involved in facilitating this unintended signal.

In summary, DNA ends play a central role in enabling artifactual probe binding. Whether via exposed overhangs or localized denaturation at termini, these features act as points of access that turn probe–DNA interactions into spurious sequencing signal.

### Comparison of CHART-seq and RAP-seq methods through the lens of Xist studies

While most studies published to date used ChIRP-seq, several mouse studies utilized the CHART-seq and RAP-seq (**Fig. 6a**). These methods were initially developed—and most frequently applied— to probe the chromatin occupancy of the *Xist* lncRNA, which plays a central role in X-chromosome inactivation in female mammalian cells, coating one of the two X chromosomes to mediate its transcriptional silencing and thereby ensuring dosage compensation for X-linked genes between males (XY) and females (XX). Because Xist is known to localize to the X chromosome, these datasets provide an opportunity to compare methodological performance across techniques for a target transcript with an established function and known genomic occupancy.

**Figure 6.**
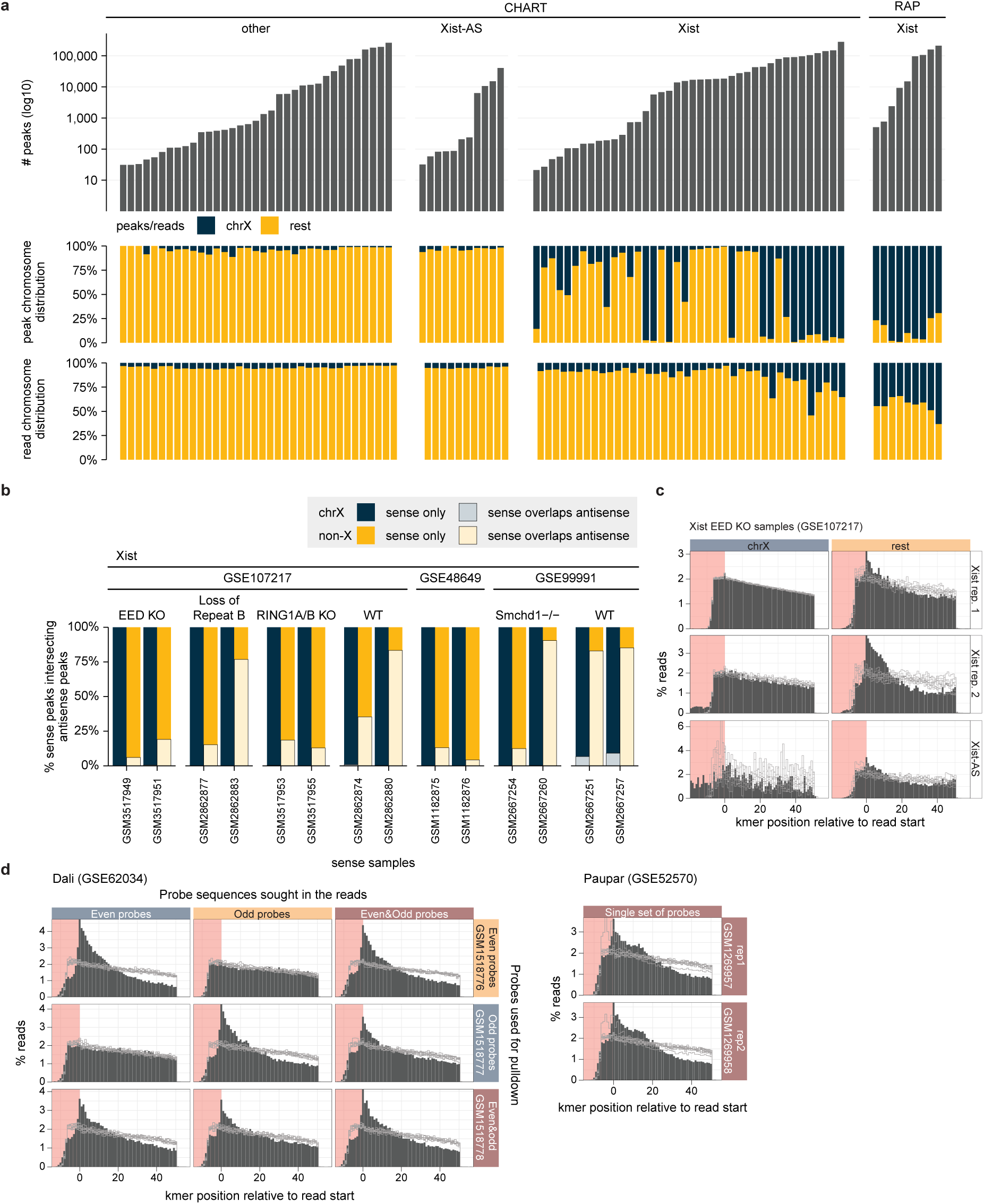
CHART- and RAP-seq method comparison. **a,** Top: Total number of peaks detected per sample, sorted in ascending order, and stratified by targeted transcript and method. Middle: Proportion of peaks assigned chromosome X or other chromosomes. Bottom: Proportion of reads assigned chromosome X or other chromosomes. **b,** Barplot of overlap between Xist CHART-seq peaks samples and their corresponding antisense probe samples peaks. **c,** Distribution of read start positions relative to the start of the longest k-mer across raw samples, stratified by chromosome X or not. **d,** For the indicated samples, distribution of read start positions relative to the start of the longest k-mer of published peaks sets across their underlying raw samples. K-mers that intersect but do not span the entire match are shaded in pink. Gray lines correspond to shuffled probes.

In RAP-seq and CHART-seq datasets probing *Xist*, we observed substantial variation in the total number of detected peaks (**Fig. 6a** top panel). Consistent with the well-established localization and function of *Xist*, a large proportion of peaks are found on the X chromosome (**Fig. 6a**, middle panel). In the RAP-seq samples, a consistently dominant fraction of peaks are on the X chromosome. In contrast, CHART-seq samples show greater variability: while many exhibit a pronounced enrichment or even a clear predominance of X chromosome peaks, others display only moderate enrichment or lack an evident bias toward the X chromosome.

Among the methods we examined, CHART-seq is the only one that employs as controls antisense probes that are reverse complement of the “on-target” probes (henceforth referred to as “antisense probes”), thus lacking in ability to hybridize to the GOI RNA, while able to bind the same DNA regions. As expected, in the *Xist* datasets, peaks obtained using antisense probes differ from the sense probes and show no detectable bias towards the X chromosome (**Fig. 6a** top middle panel). This supports the notion that the sense probe peaks are indeed RNA-driven. When comparing peaks obtained using sense vs. antisense probes, peaks on the X chromosome overlap considerably less than peaks on other chromosomes, suggesting that those non-X peaks are more likely to be driven by probe-DNA hybridization (Fig. 6b). This pattern suggests that overall, antisense probes can serve as an effective means of identifying and filtering nonspecific peaks.

When examining the overall mapping properties of the reads, RAP-seq samples consistently exhibit a high proportion (>30%) of reads aligning to the X chromosome. In contrast, most CHART-seq samples show only weak or negligible X-chromosome enrichment, both relative to antisense and to CHART-seq datasets targeting other lncRNAs (such as *Dali*, *Paupar*, B2, and *Jpx*). These findings suggest that RAP-seq captures a more specific set of *Xist*-associated fragments, resulting in overall fewer off-target or nonspecific DNA sequences compared to CHART-seq. Nevertheless, the lower read-level enrichment observed in CHART-seq does not necessarily imply reduced signal specificity at the peak level. Indeed, in several CHART-seq samples, X-chromosome enrichment becomes substantially more apparent after peak calling, indicating that although a larger proportion of reads may originate from background non-X regions, the most confidently bound regions identified by CHART-seq remain largely consistent with Xist’s established chromosomal occupancy pattern.

We then further zoomed into the CHART-seq raw read sequences and examined the position of the longest probe-derived kmer found in each peak. We found a similar skew to the read starts as was observed for many ChIRP-seq datasets (**Fig. 5e-f**). Notably, for the *Xist* studies, no distinct enrichment of read starts near the longest k-mer is observed for reads mapping to the X chromosome, while such an enrichment was seen for the non-X reads. Such signal was found in non-*Xist* CHART-seq datasets indicating considerable off-target signal (for example, **Fig. 6d**).

Overall, for the studies focused on X-inactivation, it is clear that both CHART-seq and RAP-seq are capable of enriching for real Xist-targeted regions on the X chromosome, but at least the CHART-seq studies show variability in the overall signal/noise as evidenced from variability between samples, sometimes within the same study, in the fraction of reads coming from the X chromosome and in the fraction of peaks containing probe-matching probes.

## Discussion

The central challenge in analyzing biological data is distinguishing meaningful signal from background noise. In our examination of numerous published RNA chromatin occupancy studies, we found that although they potentially hold promise for elucidating RNA-chromatin interactions, the resulting readouts are frequently compromised by fragments likely generated through partial probe hybridization, rather than through specific binding to the transcript of interest. This, in turn, leads to erroneous identification of binding sites – even when independent interleaving probe sets are used – posing a major barrier for drawing biologically meaningful conclusions from such data. We first demonstrated the issue of RNA-independent peak recovery by studying the *NESPR* lncRNA, and then extended the analysis on additional datasets, where we find presence of probe-matching *k*-mers in peaks and near 5’ ends of sequencing reads. Importantly, analyzing the *NESPR* data through the same prism shows that it has a similar enrichment of probe-matching k-mers within peaks and within reads as other published studies (**Fig. S8**).

Notably, while we find that in most studies there is clear evidence that many of the peaks result from probes enriching partially matching DNA fragments, there are several notable exceptions of studies with signal/background ratios close to one. These include several, but not all, of the *Xist* studies described above, the study of the *Haunt* lncRNA in mouse cells ^26^ and the recent study of *HOXDeRNA*^37^ in human cells. In these studies, we find no evidence for substantial enrichment of probe-derived *k*-mers in ChIRP-seq peaks (**Fig. 4d, S4b, S5**). This shows that while RNA-independent peaks are a very common problem, in certain cases it is mitigated, possibly related to the RNA abundance of the GOI or other experimental differences such as probe hybridization conditions, but those are difficult to pinpoint at the moment.

The recovery of RNA-bound DNA fragments in sequencing assays relies on the use of GOI-targeting probes. The rationale for the use of multiple probes is that multiple probes better recover the RNA of interest, as demonstrated in the study that first introduced ChIRP-seq^38^. However, unlike visualization techniques such as FISH, where a detectable signal typically arises from the combined binding of multiple probes to the same molecule, sequencing-based methods are much more sensitive and so can register a signal from a single probe. This makes sequencing approaches inherently more susceptible to artifacts from partial or nonspecific hybridization and calls for greater caution when designing sequencing-based protocols and when interpreting their results. In addition to RNase H-based probe validation, as used in CHART-seq, most studies rely on RT-qPCR analysis of the enriched RNA fraction to confirm that the GOI is specifically captured by an on-target probe set. However, small DNA fragments, such as those generated during genome fragmentation, can co-purify during RNA extraction despite DNase treatment^39^. This contamination can be relevant, as we observed that RNA-independent DNA enrichment often appears strongest at the GOI’s own genomic locus, potentially complicating the interpretation of RNA enrichment by the probe set. Depending on the genetic structure of the GOI, this can be mitigated using intron-spanning primers, which selectively amplify cDNA derived from spliced RNA rather than contaminating genomic DNA, providing a more accurate measure of RNA enrichment.

Our findings offer a framework through which RNA-chromatin interaction studies can be improved and refined through experimental or bioinformatic adjustments. In particular, the characteristic pattern of reads intersecting or flanking genomic matches of probe *k*-mers, suggests a basis for potential solutions. Given that DNA ends appear particularly susceptible to probe binding, minimizing or eliminating staggered ends may help reduce, or even eliminate, noise arising from probe-to-DNA interactions. This bias could potentially be addressed by fragmenting the DNA using a cocktail of blunt-end restriction enzymes, as employed in other sequencing protocols, or by enzymatically blunting DNA ends prior to probe hybridization. In addition, as demonstrated for *NESPR*, incorporating multiple negative controls such as RNase-treated conditions or a cell line that is not expressing the GOI in the experimental design, can strengthen confidence in distinguishing true RNA-chromatin interactions from RNA-independent DNA enrichment. Also, as we show in the *Xist* CHART-seq studies, the rarely used antisense probes are particularly useful for identifying peaks likely resulting from probe-DNA hybridization.

Notably, several protein-DNA sequencing methods have recently shifted from *in vitro* pulldown strategies, such as ChIP-seq, to *in situ* target-fragmentation approaches, including ChEC-seq, CUT&RUN, and CUT&Tag^40–42^. Whereas the former relies on the differential enrichment of fragments of interest from a background of all fragments, the latter generates and captures DNA fragments only at genuine binding events, leaving most of the genome intact. Adapting RNA-to-DNA mapping techniques to follow a similar *in situ* paradigm may help eliminate non-specific probe-to-DNA binding, thereby enhancing signal specificity and clarity, while also likely reducing the amount of required material/cells. Notably, several recent methods describe methodologies based on proximity labeling described RNA-chromatin interactions. These include O-Map that was used for mapping chromatin occupancy of Xist^43^, and related methods were used for profiling proteins and RNAs, but not yet DNA, near RNAs of interest^44,45^. A potential challenge in using these approaches for RNA-chromatin interaction mapping is the difficulty in extrapolating between the enrichment of a specific region of interest to the actual physical distance between the RNA and the genomic region, and/or the frequency of the interactions.

On the bioinformatic side, we provide a heuristic framework for identifying problematic reads or regions. We have shown that probe-derived *k*-mers tend to appear disproportionately near the beginning of reads—a property that can be leveraged to filter out *k*-mer matches of length ≥7 that occur within the first 20 nucleotides of a read. However, this approach has a notable limitation: it does not remove reads that flank the *k*-mer on its 3′ side as *k*-mer sequences that led to the fragment enrichment may have been removed during blunting in library preparation. Our analysis further reveals that reads often begin near the *k*-mer site, whether directly intersecting or flanking it. This suggests that problematic regions could also be identified as short genomic windows with a high density of read starts relative to overall coverage. Another, potentially bigger challenge is the difficulty to determine a particular threshold above which a peak or a read can be flagged as potentially problematic. For example, when considering peaks, the strongest signal-to-noise ratio is often achieved for *k*~12, but removing just regions with matches of this length will likely retain many spuriously recovered regions containing just shorter matches, or imperfect probe homology with mismatches. This limitation makes it difficult at present to provide informatically a “clean” list of confident peaks for the available studies.

Among the different methods for chromatin-RNA interaction profiling, ChIRP-seq has been much more extensively adopted than CHART-seq and RAP-seq, likely because of the relative ease of probe design, without the RNase H-based pre-screening recommended for CHART-seq, and the low costs of synthesis of short probes. The other two methods were mostly used in *Xist* studies (and so the same optimized probes could be reused). For CHART-seq we do see that in some studies there is evidence for probe-based peak recovery, and so it does not appear immune to the problems our analysis highlights. Because CHART-seq and RAP-seq were mostly used to profile *Xist*, which is also the RNA with the best evidence for occupying chromatin distally from its site of transcription, it is difficult for us to effectively compare the different protocols. What appears certain is that, when ChIRP-seq is employed, it is important to at least use of even and odd probe sets, carefully study the consider the overlap in the recovered peak, and to examine probe-derived *k*-mer enrichments. More importantly, it is crucial to both perform controls, particularly ones in which the RNA of interest is depleted or absent and the experiment is repeated, as we demonstrate here with *NESPR*.

LncRNAs as a group were suggested over a decade ago to often act as ‘traffic controllers’ assisting chromatin-associated protein complexes with no apparent sequence specificity, such as PRC2, to reach specific distal sites^46^. This notion, if prevalent, has two prerequisites – first, that the RNAs can specifically recognize chromatin-related protein complexes, and the second that the RNAs can, independently, identify specific distal sites. The first part of this dogma has been recently cast in doubt^47–51^, although it remains contested at least for PRC2^52,53^. Our result here adds concern to the second prerequisite of the dogma, as we show that the evidence for widespread specific lncRNA-chromatin interactions remains weak. In many among the dozens of studies we analyzed here, probe-based DNA recovery appears as a major issue that was not tested or addressed. Typically, the specific RNA-chromatin target mapping was not the main experiment performed, yet it was typically the major evidence that the RNA (mostly lncRNA) is recruited to specific distal regions. Based on our results, at present time, it is difficult for us to determine if most or all of the distal binding sites reported in most studies originate from probe hybridization to exposed DNA ends. Importantly, validation by qPCR, which is employed in some of the studies, is prone to the same concerns as the sequencing-based approaches. Therefore, generally, we can cast substantial doubt on the notion that many lncRNAs are indeed interacting specifically with distal loci. This doubt resonates with the typically low copy number of most lncRNAs which would make it difficult for them to effectively reach and regulate the thousands of sites they are often reported to occupy. Even when accounting for the possibility of temporal DNA binding by the RNA of the GOI, it remains hard to reconcile how a low abundance lncRNA could engage genomic sites whose number exceeds its copy number by orders of magnitude. Moreover, our results may have implications for ChIRP-, CHART-, or RAP-seq-based strategies to identify RNA-protein interactions, such as ChIRP-MS, where DNA is not routinely depleted. Enriched proteins may originate from RNA-independent DNA fragments, carrying little to no biological relevance for the molecular mechanism or life cycle of the GOI.

It remains to be determined whether modifications to the protocol, study design, or analysis pipeline can significantly improve RNA chromatin occupancy sequencing methods results. Until such improvements are validated, our results show this data should be interpreted with caution, and conclusions drawn from it should be considered as source for hypothesis generation rather than as validation for targeting of specific regions by RNA molecules.

## Material and methods

### Cell culture

IMR-32 and SHEP cell lines were cultured in RPMI 1640 supplemented with 10% FBS, 2 mM L-glutamine, and 30 U/mL Penicillin-Streptomycin. HEK293T cells were cultured in Dulbecco’s Modified Eagle Medium (DMEM) supplemented with 10% FBS, 1% sodium pyruvate, and 1% GlutaMAX. Cells were grown at 37°C at 5% CO2. Cells were maintained at a maximal confluence of 50-60% and passaged twice a week. Cells were frequently monitored to be mycoplasma-free using a mycoplasma PCR detection kit.

### Lentiviral production and transduction of shNESPR

#### Cloning of shRNAs

Short hairpin RNAs targeting *NESPR* exonic sequences were designed based on LNCASO software (https://iomics.ugent.be/lncaso/; sequences in **Table S2**). Scrambled shRNA sequence was derived from Cai *et al.* (2006). Oligonucleotides comprising the top and bottom of the shRNA contained AgeI and EcoRI overhangs, respectively. Top and bottom oligonucleotides were phosphorylated using T4 Polynucleotide kinase (NEB) according to the manufacturer’s instructions, and subsequently annealed by adding 50 mM NaCl and ramping down the reaction from 100°C to 4°C with 0.1°C/s steps in a thermocycler. Oligoduplexes were ligated in AgeI/EcoRI-digested pLKO.1 vector (Addgene #8453) using 1 U T4 DNA ligase (ThermoFisher Scientific). Ligation mixtures were chemically transformed in DH10B competent cells, and grown on LB agar plates containing 50 μg/mL carbenicillin. Colonies were verified by restriction digest and Sanger sequencing.

#### Lentiviral production and transduction

6.5 × 10^6^ HEK293T cells were seeded for calcium phosphate transfection the next day. Medium was refreshed 30 min prior to transfection. The DNA mixture was comprised of 24 μg of shRNA transfer plasmid, 18 μg pCMV-dR8.74, 7.2 μg pMD2.G, and 75 μL of 2.5 M CaCl_2_ in a total volume of 750 μL in cell culture-grade water. The transfection mixture was added dropwise to 750 μL 2X BioUltra HEPES-buffered saline solution (Sigma-Aldrich) while vortexing. This transfection mixture was incubated for 5 min at room temperature, and added to the cells. On the next day, a Ca_3_(PO_4_)_2_ precipitate was observed and the medium was refreshed to prevent cytotoxicity due to the transfection reagents. The following two days, culture medium containing lentiviral particles was collected and kept at 4°C. Three days after transfection, both harvests were pooled and spinned at 500 x g for 5 min at 4°C to pellet debris. Supernatant was filtered through a 0.45 μm filter, and transferred to an ultracentrifugation tube. Lentiviral particles in the supernatant were pelleted by ultracentrifugation for 2 h and 10 min at 100,000 x g at 4°C. Lentiviral pellets were resuspended in 100 μL culture medium, and stored at −80°C. A colony formation assay was performed with 2 μL concentrated virus to determine lentiviral titers. For lentiviral transductions, 1 × 10^6^ IMR-32 cells were seeded. The day after seeding, concentrated lentivirus was added to achieve an MOI of 10. The next day, the culture medium was refreshed. Two days after transduction, 1 μg/μL puromycin was supplemented to the culture medium to select for shRNA-transduced cells. After initial selection, cells were maintained in puromycin-free cell culture medium. One week prior to any shRNA experiment, cells were spiked with 1 μg/μL puromycin to evaluate transgene silencing.

### RNA extraction, cDNA synthesis and RT-qPCR

#### RNA extraction

Cells were washed once with PBS, trypsinized, and collected by centrifugation. After two additional PBS washes, pellets were resuspended in 700 μL QIAzol, incubated for 15 min at room temperature, snap-frozen in liquid nitrogen, and stored at −80°C until further processing. Samples were thawed at room temperature, and 140 μL 100% chloroform was added for phase separation. After vigorous mixing and centrifugation at 12,000 x g for 15 min at 4°C, the upper aqueous phase was collected to which 600 μL 100% EtOH was added. RNA from the mixture was isolated with the miRNeasy Micro kit (Qiagen) according to the manufacturer’s instructions, including a DNase I digest. RNA concentrations were measured on a nanodrop, and RNA was stored at −80°C.

#### cDNA synthesis

Complementary DNA synthesis was performed with the PrimeScript RT kit (Takara Bio) or iScript Advanced cDNA synthesis kit (Bio-Rad) following the manufacturers’ instructions with RNA input up to a maximum of 500 ng RNA template with maximal shared RNA input across samples where required. For validating ChIRP *NESPR* enrichment, a maximal volume was used in the reaction for all samples.

#### RT-qPCR of NESPR after ChIRP enrichment

*NESPR* was amplified using primers spanning its intron. All primers are listed in **Table S4**. Quantitative PCR reactions were performed in 10 μL total volume containing 5 μL 2X SsoAdvanced Universal SYBR Green master mix, 0.5 μL of 5 μM forward and reverse primer each, and 4 μL 4X diluted cDNA in nuclease-free water. Samples were measured in technical quadruplicates.

Fluorescent signal was measured on a LightCycler 480 system using the following cycling conditions: 1 cycle at 95°C for 5 min, 40 cycles at 95°C for 10 s, 60°C for 10 s, and 72°C for 10 s, followed by melting curve analysis. Quantitation cycles (Cq) were normalized to GAPDH. Normalized Cq (ΔCq) were compared relative to the input sample to calculate *NESPR* fold enrichment in each sample.

### Next generation sequencing

#### RNA-sequencing and data processing

Experiments contained four biological replicates per shRNA, with each replicate transduced independently at an MOI of 10. RNA libraries were prepared from 8 μL purified RNA of shRNA-transduced IMR-32 cells using the TruSeq Stranded mRNA Library Prep Kit LT according to the manufacturer’s instructions. RNA was fragmented at 94°C for 4 min prior to first strand cDNA synthesis. Library amplification was performed with 12 PCR cycles. Library size analysis was assessed on fragment analyzer. Library quantification was performed on a Qubit 4 Fluorometer. TruSeq libraries were pooled, the pool was quantified by Qubit, and 2 pM with 1% PhiX was loaded on a NextSeq 500 instrument (NextSeq 500/550 High-Output kit, 75 cycles, single-end sequencing). Reads were quality controlled using FastQC (v.0.11.8) and trimmed with cutadapt (v1.18) to remove any low quality bases and Illumina adapter sequences. Trimmed reads were aligned with Bowtie 2 (v2.3.2) to the human hg19 reference genome (GRCh37) followed by quantification using HTSeq (v0.6.1). Count data was imported in R, and differential analysis was performed with DESeq2 (v1.42.0) with shRNA as the only design variable (design = ~ shRNA). PCA confirmed replicate clustering per conditions (shSCR and shNESPR). DESeq2-normalized counts of shNESPR were contrasted to shSCR, with log2 transformation of the resulting fold changes per gene. Statistical significance was determined using the default DESeq2-implemented Wald test with a 5% adjusted P-value cutoff to support significance. Multiple testing correction was performed with Benjamini-Hochberg. Genes with a baseMean of less than 10 were removed.

#### ChIRP-sequencing and data processing

ChIRP-sequencing protocol was derived from Chu *et al.* (2011) and Chu *et al.* (2012). In brief, 20-30 million IMR-32 or SHEP cells per sample were cultured in T175 flasks and harvested by scraping in ice-cold PBS. *In cellulo* cross-linking was done by resuspending the cell pellets in 10 mL 1% glutaraldehyde in PBS followed by end-over-end rotation for 10 min at room temperature. The reaction was quenched by addition of 1 mL 1.25 M glycine and incubation for 5 min at room temperature with end-over-end rotation. Crosslinked cells were pelleted by centrifugation and resuspended in 2 mL SDS lysis buffer (50 mM Tris-HCl pH 7.5, 10 mM EDTA, and 1% SDS) supplemented with 60 U/mL SuperaseIn RNase inhibitor, 1 mM DTT, 0.5 mM PMSF, and cOmplete Protease Inhibitor Cocktail. Lysates were sonicated with a Bioruptor Pico sonication device cooled to 4°C using 30 s ON/30 s OFF cycles until the lysate was cleared. Intermittent samples were taken for fragment analysis, and sonication was continued until genome fragmentation to 500 bp. The sonicated cell lysate was centrifuged at 16,100 x g for 10 min at 4°C, and the supernatant was transferred to a DNA LoBind microcentrifuge tube. From these lysates, 10% was taken as an RNA input sample, and 1% as a DNA input sample. For RNase treatment, 10 μL of 10 μg/μL RNase A and 5 μL of 10 U/μL RNase H was spiked into the lysate, and incubated for 30 min at 37°C. Per sample, 100 μL Dynbeads MyOne Streptavidin C1 were rendered RNase-free following the manufacturer’s instructions, and were conjugated with 625 pmol of 3’-biotinylated *NESPR*- or *lacZ*-targeting raPOOL probes (siTOOLs Biotech; probe sequences are listed in **Table S4**). Lysates were pre-cleared with 30 μL equilibrated non-conjugated beads for 30 min at 4°C. Pre-cleared lysates were incubated with probe-conjugated beads for 3 h at 4°C with vertical mixing. Beads were washed three times with unsupplemented SDS lysis buffer, and resuspended in 1 mL unsupplemented SDS lysis buffer. Ten percent of the bead suspension was taken for RNA isolation to confirm RNA enrichment, and 90% was used for DNA isolation and downstream processing. DNA fractions were magnetized, and the beads were resuspended in 150 μL DNA elution buffer (50 mM NaHCO3 and 1% SDS). DNA was eluted in two consecutive rounds by incubation with 1 μL of 10 μg/μL RNase A and 1 μL of 10 U/μL RNase H. Next, supernatant was digested with 1.5 uL of 800 U/mL proteinase K at 50°C for 45 min, and DNA was purified using 5PRIME Phase Lock Gel Heavy (Quantabio) and PhOH:Chloroform:Isoamyl following the manufacturer’s instructions. The aqueous phase was transferred, and 3 μg GlycoBlue, 30 μL of 3 M NaOAc, and 900 μL of 100% EtOH was added and stored overnight at −20°C. On the next day, samples were spinned at 16,100 x g for 30 min at 4°C. Supernatant was removed and the pellet was washed once with 1 mL 70% EtOH. The pellet was air-dried and resuspended in 30 μL nuclease-free water.

ChIRP DNA was size-selected on 2% E-gel to remove GlycoBlue carrier. DNA fragments between 200-650 bp were excised, resuspended in 650 μL Agarose Dissolving Buffer, and purified using the Zymoclean Gel DNA Recovery kit (Zymo Research). DNA was eluted in 52 μL nuclease-free water. DNA concentration was determined using Qubit, and 50 μL of each sample was used as input for library preparation with the NEBNext Ultra II DNA Library Prep kit following the manufacturer’s instructions with NEBNext adaptors. After adaptor ligation, 0.2 U/μL USER enzyme was added to remove uracil bases. Libraries were purified with AMPure XP beads at a 0.9X ratio and eluted in 40 μL nuclease-free water. Input libraries were amplified for 9 cycles, whereas libraries of the enriched fractions were amplified for 18 cycles using 20 μL input. Amplified libraries were clean with AMPure XP beads at a 0.8X ratio and eluted in nuclease-free water. Library fragment distribution were assessed using a fragment analyzer. Concentrations were quantified using the KAPA Library Quantification kit, after which libraries were equimolarly pooled at a final concentration of 1.94 nM. A final loading concentration of 650 pM was used for sequencing on a NextSeq2000 instrument (NextSeq2000 P2, 100 cycli, single-end sequencing), including 2% PhiX for sequencing QC purposes. Raw sequencing data were evaluated using FastQC (v.0.11.8) and subsequently mapped to the hg19 human reference genome (GRCh37) with Bowtie2 (v2.3.2). Peak calling was performed using MACS2 (v2.1.0) only retaining peaks that passed the 0.05 q-value threshold. Peaks in IMR-32 samples were filtered for chr2p amplified regions.

### Distance of *NESPR* ChIRP peaks to DEGs and permutation testing

DEGs were stratified by absolute log2FC (|log2FC| ≥ 0.5 or ≥ 1) into moderate and strong DEGs. The TSS of each DEG was obtained using biomaRt (v.2.58.2) and converted to genomic ranges with GenomicRanges (v.1.54.1). To quantify the fraction of DEGs near *NESPR* peaks, the proportion of DEGs with at least one ChIRP peak within 10 kb or 50 kb of the TSS was calculated using distanceToNearest as implemented in the GenomicRanges package. Statistical significance was assessed by permutation testing using regioneR (v.1.34.0), performing 1,000 randomizations of the ChIRP-seq peak positions across the hg19 genome. Randomized peaks preserved chromosome distribution and overall width distribution of the original peak set.

### RNAscope

10,000 IMR-32 or SHEP cells were seeded in 400 μL medium per chamber of an 8-chamber imaging slide and grown to 70-80% confluency. On the next day, half the medium was replaced with 4% pre-warmed PFA in PBS for a final concentration of 2% PFA. After 15 min incubation at room temperature, the total volume was replaced with 2% pre-warmed PFA in PBS, and incubated for an additional 10 min. Slides were washed three times with PBS. RNAscope chromogenic staining was performed with custom NESPR-specific probes using the RNAscope 2.5 HD reagent Kit-RED (Advanced Cell Diagnostics) according to the manufacturer’s instructions.

### Obtaining data from published studies

Consensus peak sets from studies were collected from the Gene Expression Omnibus (GEO), published articles, and directly from corresponding authors. Probe sequences were curated from published papers and GEO records. To ensure consistency across datasets, probe names were reformatted to a uniform style, which may differ from the original designations used in the primary publications. For datasets reported in genome assemblies differing from those used here, coordinates were converted using liftOver from Kent utilities ^54^. The number of successfully lifted peaks is provided in **Table S3** (see “Notes” column). Target transcript coordinates (**Table S5**) were inferred based on the best available evidence. When transcript IDs were provided, genomic coordinates were directly extracted. In other cases, coordinates were inferred from supporting information, including published figures, probe sequences, FISH probe data, qRT-PCR primers, and RACE experiments. These sources were collectively used to determine the most accurate genomic representation of each target transcript.

### Processing of published datasets

Raw sequencing data were downloaded from the Sequence Read Archive (SRA) and mapped to their respective reference genomes using Bowtie2 ^55^. Human samples were aligned to the hg19 reference genome (GRCh37) and mouse samples to the mm10 reference genome. A complete list of analyzed samples is provided in **Table S1**. Reads aligning to the ENCODE blacklist (https://github.com/Boyle-Lab/Blacklist/tree/master/lists) were removed prior to peak calling to avoid spurious signal at regions known to display artifactually high background in pull-down experiments ^20^. Peak calling was performed using MACS2 ^18^.

### Classification of Common and Very Common Peaks

Peaks from individual samples were merged to generate a comprehensive reference peak set. Samples targeting (i) multicopy elements (e.g., 7SK, ERV-9) or (ii) transcripts probed in multiple independent studies were excluded to avoid biasing the merged peak set. Each individual sample’s peaks were intersected with the merged set, and for every merged peak, the number of intersecting samples and studies was counted. Merged peaks overlapping peaks from ≥10 studies were defined as “very common peaks”, while those overlapping peaks from ≥6 studies were defined as “common peaks” (**Tables S6-S10**).

### Intersection and Union Analyses of Matched Probe Sets

For studies employing “even” and “odd” probe sets, samples were paired according to cell line, treatment, and genotype. Peak intersections and unions between paired samples were computed using bedtools intersect.

Expected random overlap between pairs was estimated by randomly placing, separately for each chromosome, the peaks, and counting the number of overlaps.

### Peak Processing for Sequence Analyses

Individual sample peaks, paired-sample intersections and unions, and published peak sets were analyzed for k-mer occurrence and motif enrichment.

#### k-mer Occurrence Analysis

Probe-derived *k*-mers ranging from 7 to 20 nucleotides were searched across peaks using a custom script. As controls, ten dinucleotide-shuffled variants were generated for each probe and analyzed in parallel. When overlapping matches of length *k*–1 were detected, only the first *k*-mer match was recorded (For example, if two 7-mer matches overlap by six nucleotides, the overlapping region forms a single continuous sequence that is equivalent to an 8-mer match).

#### k-mer signal-over-background

For each sample, the fraction of peaks containing an actual probe-derived *k*-mer was divided by the mean fraction of peaks containing a *k*-mer matching a shuffled probe (adding a 0.001 pseudocount to prevent division by zero). For paired samples, the mean signal-over-background value between “even” and “odd” probe sets was computed. For samples with only a single probe set, the mean value across all samples with identical experimental conditions was used.

#### Longest k-mer analysis

The longest k-mer identified within each peak was determined using the output of the *k*-mer counting script. In cases where the same longest *k*-mer appeared multiple times, all occurrences were retained.

#### Analysis of reads intersecting or flanking the longest k-mers

Because some control shuffle probes’ longest *k*-mer matches overlapped with those of the true probes, potentially confounding signal interpretation, shuffled-probe *k*-mers overlapping true-probe *k*-mers by ≥7 bases were excluded.

Reads intersecting the longest k-mer matches of both true and control probes were extracted, and duplicates were removed using MarkDuplicates from Picard (https://broadinstitute.github.io/picard/). The position of each *k*-mer relative to the read start was then determined. Similarly, reads flanking— but not overlapping—the longest *k*-mer matches of true and control probes were extracted and processed.

### Motif Discovery and Alignment to Target Transcripts

Motif enrichment analyses were performed using STREME ^56^ and HOMER ^57^. To reduce computational load, analyses were restricted to 10,000 peaks per dataset. For individual samples, the top 10,000 peaks were selected based on the “score” column of the MACS2 narrowPeak output (calculated as –log10(qValue) × 10). For published consensus peaks, where ranking information was available, the top 10,000 were used; otherwise, 10,000 peaks were randomly sampled using *bedtools sample* ^58^. All peaks were used for the intersection and union analyses of paired “even” and “odd” probe sets. STREME was run with default parameters. HOMER was executed using *findMotifsGenome.pl* to search for motifs ranging from 8 to 15 nucleotides—the same default range explored by STREME—within a 200 bp window (default). While STREME automatically optimizes motif width within the specified range, HOMER requires explicit user-defined motif lengths, which can lead to shorter motifs matching and contained within longer reported motifs.

Identified motifs were mapped back to their respective target transcripts using FIMO ^59^ from the MEME suite with default settings.

## Data availability

The ChIRP-sequencing data generated in this study have been deposited to NCBI’s Gene Expression Omnibus (accession number: GSE307444). All data will be made public upon publication.

## Funding

This work was supported by Fonds Wetenschappelijk Onderzoek (1275923N to L.D., G0D8619N to P.M.), Bijzonder Onderzoeksfond Universiteit Gent (BOF22/PDO/024 to L.D., BOF16/GOA/023 and BOF.GOA.2022.003.03 to F.S. and S.E.), Stichting Tegen Kanker (FAF-F/2018/1176 to P.M.), and the European Research Council (Consolidator Grant lncIMPACT to I.U.).

## Acknowledgement

The authors acknowledge the NXTGNT Ghent University sequencing core facility and ERA Platform Research Core (VUB) for their assistance on the presented work. We thank Noam Stern-Ginossar, Schraga Schwartz and members of the Ulitsky lab for helpful discussions and comments on the manuscript.

## Competing interests

The authors declare no competing interests.

## Supplementary Tables

**Table S1.** Processed published studies, including information on the methods, probes and their types and different treatments and controls.

**Table S2.** Probes used in the different studies, including for NESPR.

**Table S3.** Published peak sets reported in the different studies.

**Table S4**. Primers and probes used for studying NESPR.

**Table S5**. GOI coordinates.

**Table S6.** Very common peaks in the human datasets.

**Table S7.** Common peaks in the human datasets.

**Table S8.** Very common peaks in the mouse datasets.

**Table S9.** Common peaks in the mouse datasets.

## Notes

### Competing Interest Statement

The authors have declared no competing interest.

