## Supplemental Figures for "Widespread DNA off-targeting confounds studies of RNA chromatin occupancy"

### Supplementary Figures for "Widespread DNA off-targeting confounds studies of RNA chromatin occupancy" by Goldrich, Dalhaye et al.

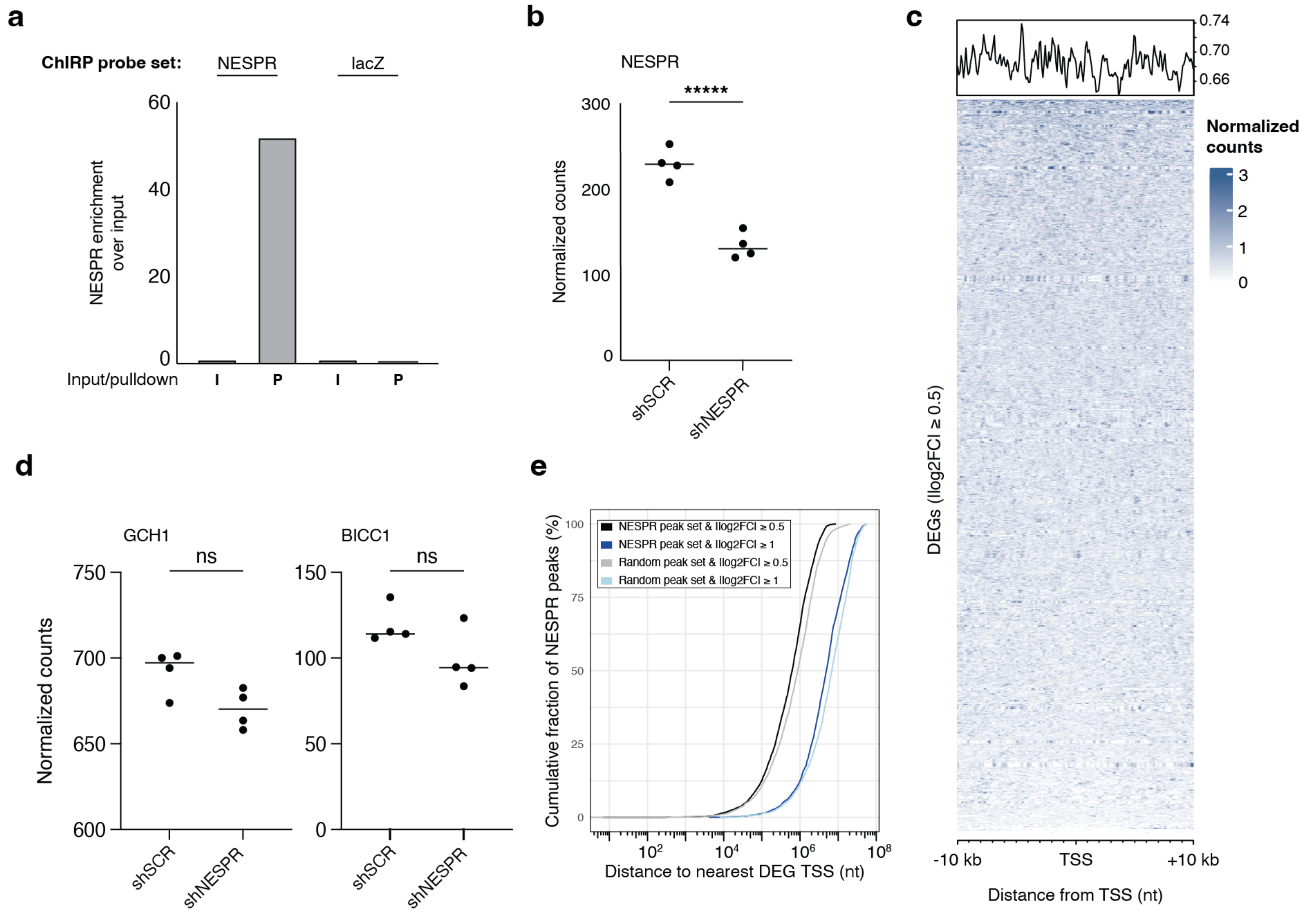


**Figure S1. *NESPR* ChIRP-seq peaks are not linked to gene expression changes upon *NESPR* depletion.** **a,** Normalized *NESPR* enrichment over input in ChIRP-seq experiments using *NESPR* and *lacZ* capture probe sets. **b,** *NESPR* expression levels in shNESPR and shSCR IMR-32 cell lines. **c,** NESPR ChIRP-seq coverage at DEGs after *NESPR* knock-down. d, Expression levels of *GCH1* and *BICC1* genes with a promoter-proximal *NESPR* ChIRP-seq peak. **e,** Distance of *NESPR* ChIRP-seq peaks to the TSS of DEGs after *NESPR* knock-down. I, input; P, pulldown; DEG, differentially expressed genes; nt, nucleotides.

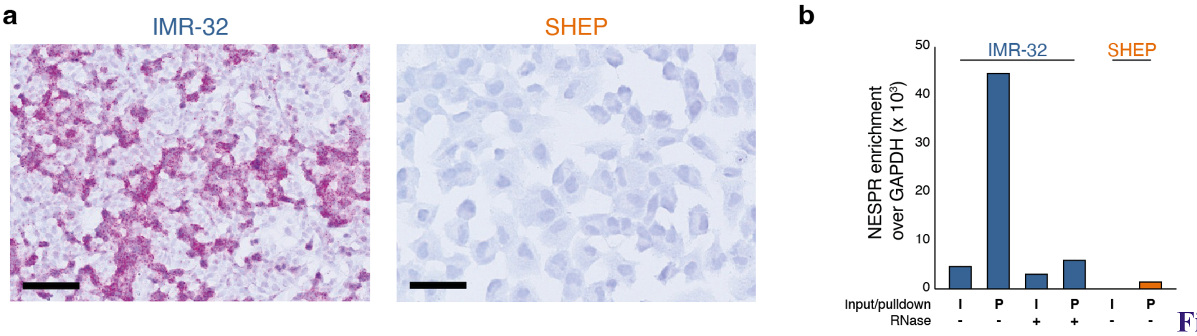


Figure

**Figure S2. NESPR expression and *NESPR* enrichment in IMR-32 and SHEP cells. a,** Colorimetric RNAscope analysis of *NESPR* in IMR-32 and SHEP cells. **b,** *NESPR* enrichment over GAPDH in ChIRP-seq fractions in IMR-32 (with and without RNase treatment) and SHEP. Dark blue, cells with *NESPR* expression; orange, cells without *NESPR* expression. Scale bar shows 20 µm.

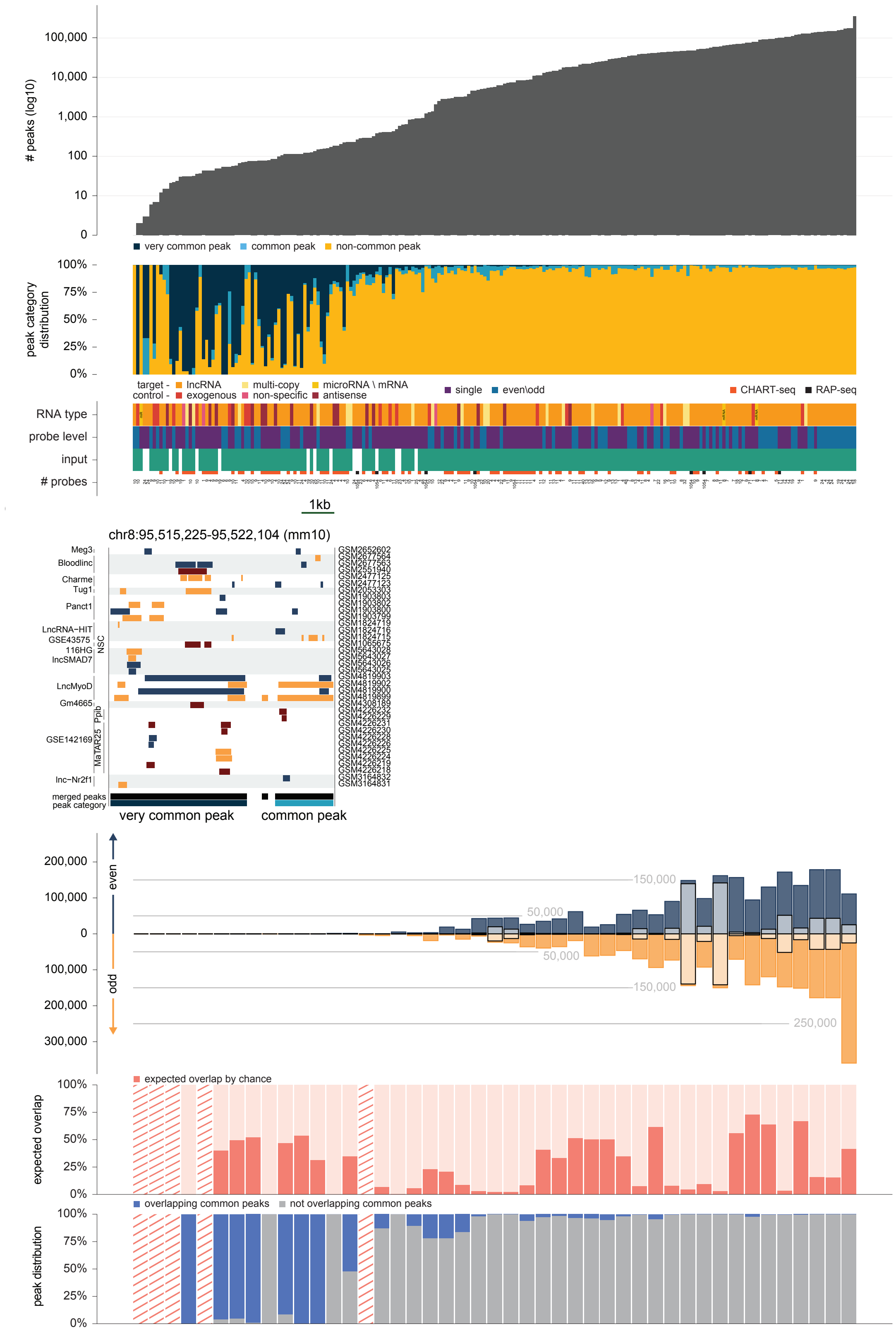


**Figure S3. Peak distributions and properties across mouse samples, including matched replicates. a,** As in Fig. 3b for mouse samples. **b,** An example of a region in the mouse genome containing a very common peak and a common peak, with the peaks from the individual studies overlapping it. **c,** As in Fig. 3c for mouse samples.

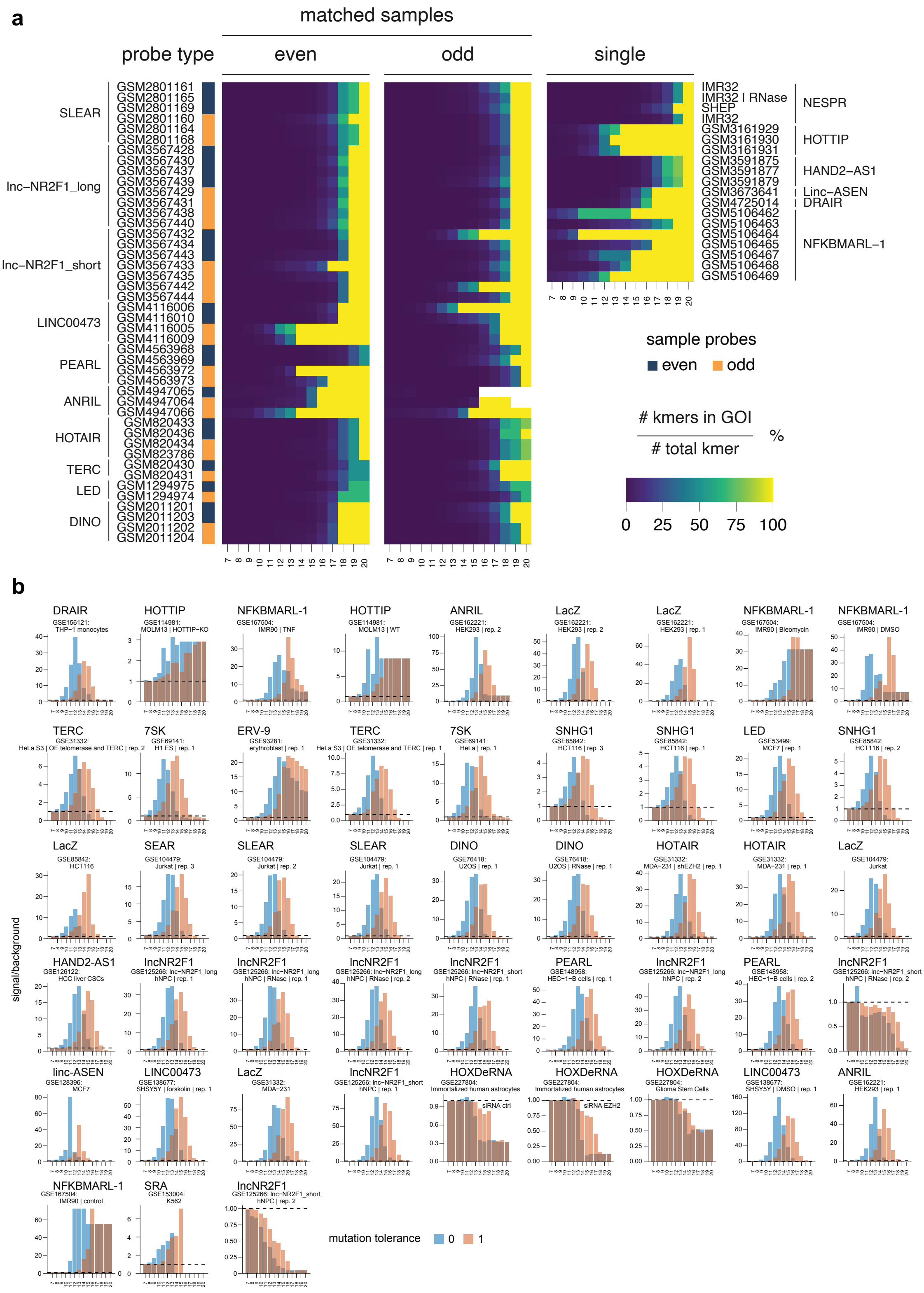


**Figure S4. Signal-over-background profiles of peak k-mers in human samples.**

a) Of the peaks that have a kmer match of the indicated length, the color indicates that fraction that occured in peaks that overlap the GOI. b)Signal-over-background values are plotted for k-mers of length 7 to 20 across grouped human samples. Sample groups are ordered by the average number of peaks per group, arranged from left to right and top to bottom. Groupings were defined based on identical experimental conditions, probe target, and biological replicate number. When applicable, both “even” and “odd” probe sets were included; otherwise, single-probe sets (“single”) were used.

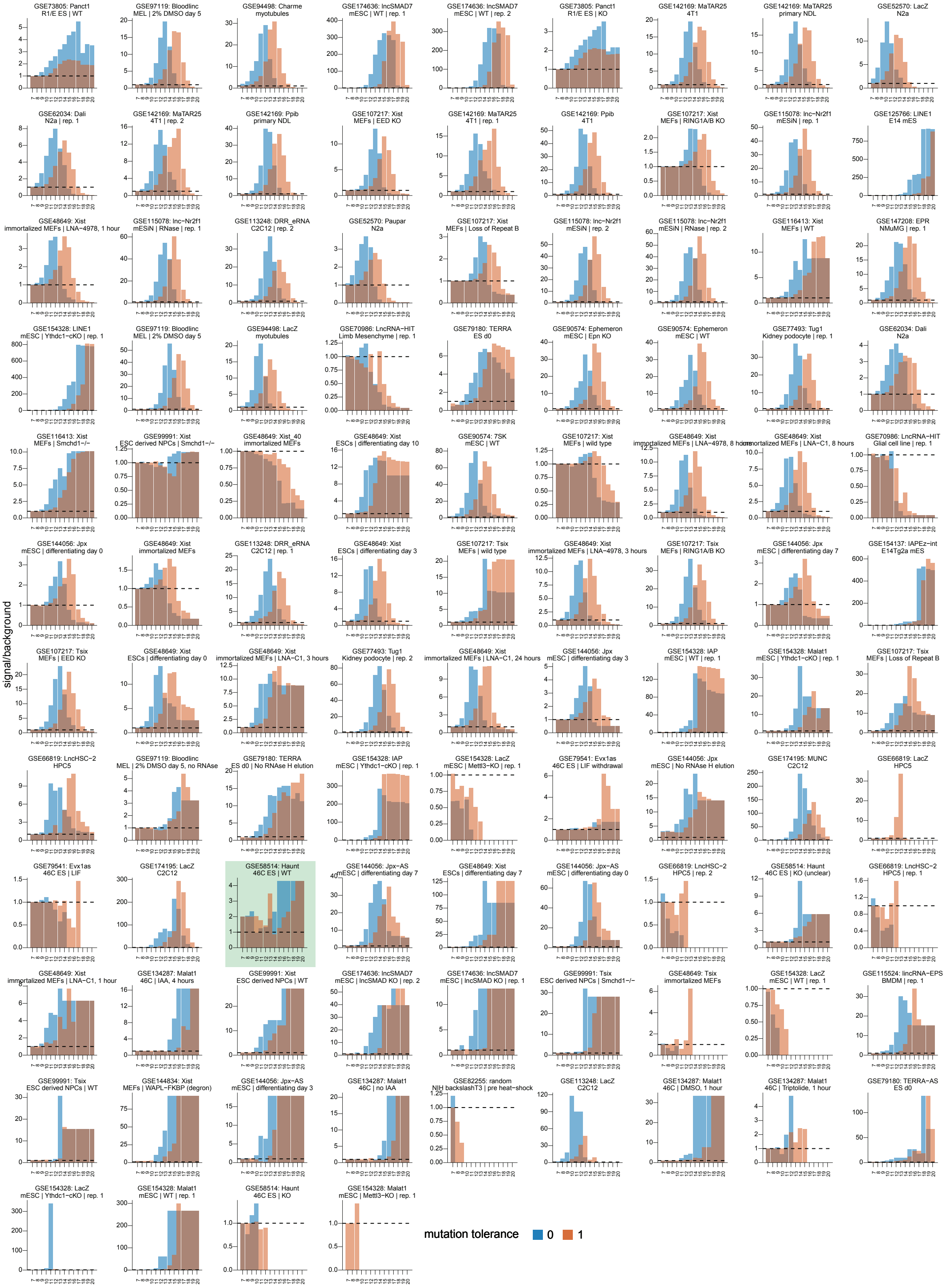


**Figure S5. Signal-over-background profiles of peak k-mers in mouse samples.** Signal-over-background values are plotted for k-mers of length 7 to 20 across grouped mouse samples. Sample groups are ordered by the average number of peaks per group, arranged from left to right and top to bottom. Groupings were defined based on identical experimental conditions, probe target, and biological replicate number. When applicable, both “even” and “odd” probe sets were included; otherwise, single-probe sets (“single”) were used. The Haunt lncRNA study [23] mentioned in Discussion is shaded.

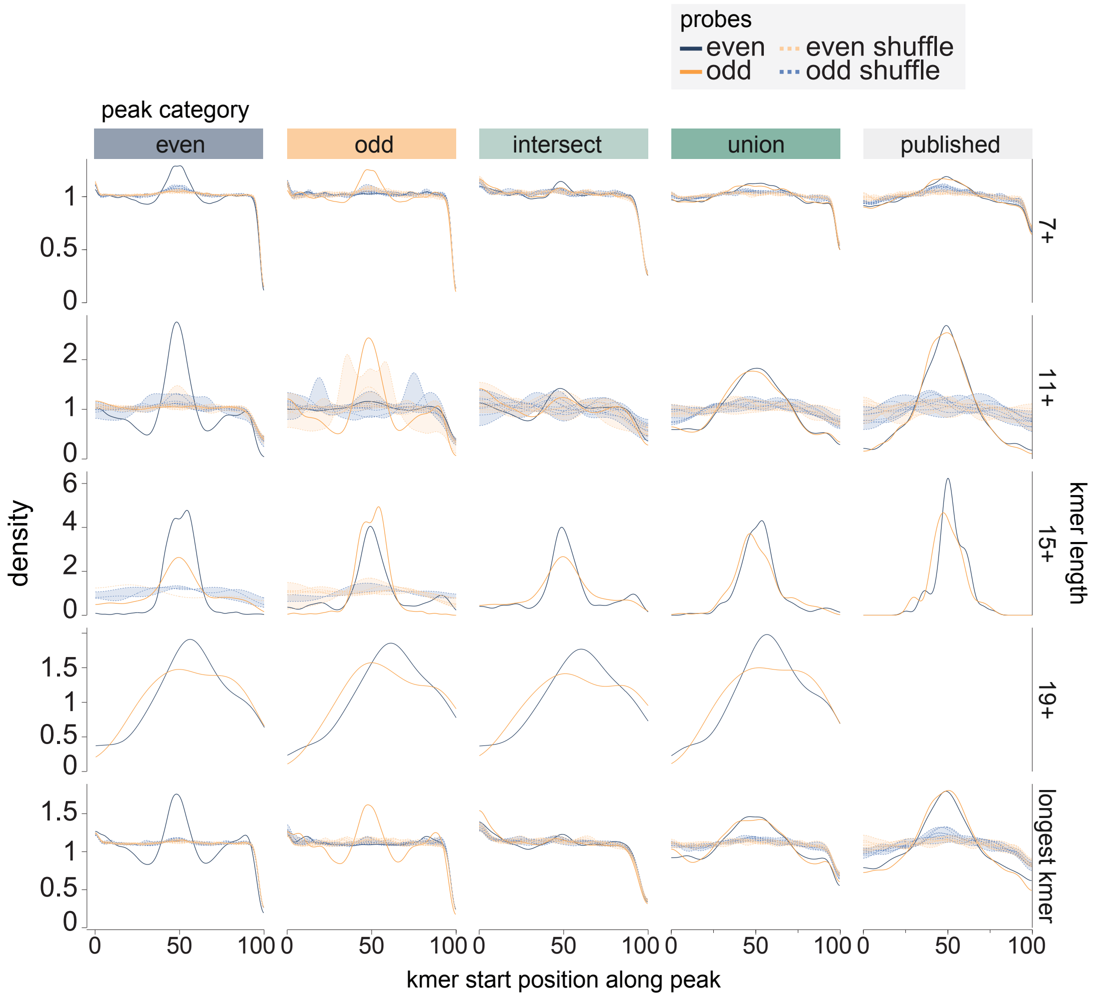


**Figure S6. Relative positions of k-mer hit starts across peak sets in a dataset for the LED lncRNA.** Distributions of the start positions of true (bold lines) and shuffled (dashed lines) k-mer hits are shown across different peak sets, for k-mers of length ≥7, ≥11, ≥15, ≥19, and for the longest k-mer per peak. Only probe peak sets with at least 25 peaks are shown to avoid plotting potentially spurious distributions.

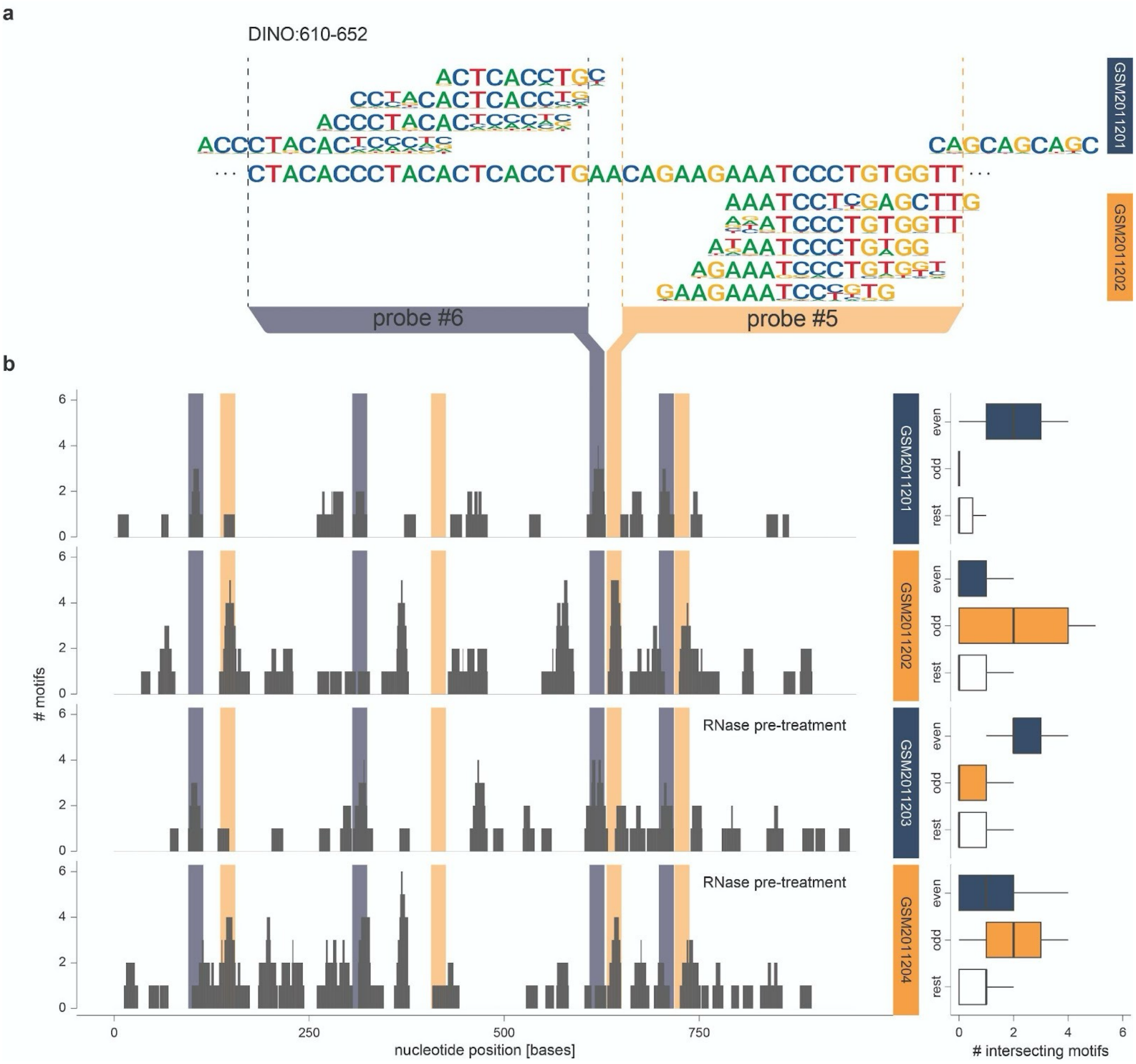


**Figure S7. Motif coverage along the *DINO* lncRNA transcript. a,** Aligned motifs along "even" and "odd" probe regions in *DINO* transcript. **b,** Left: Distribution of motif coverage along the *DINO* lncRNA transcript. Right: Boxplots of motif coverage stratified by whether the nucleotide overlaps with probe target sites or not.

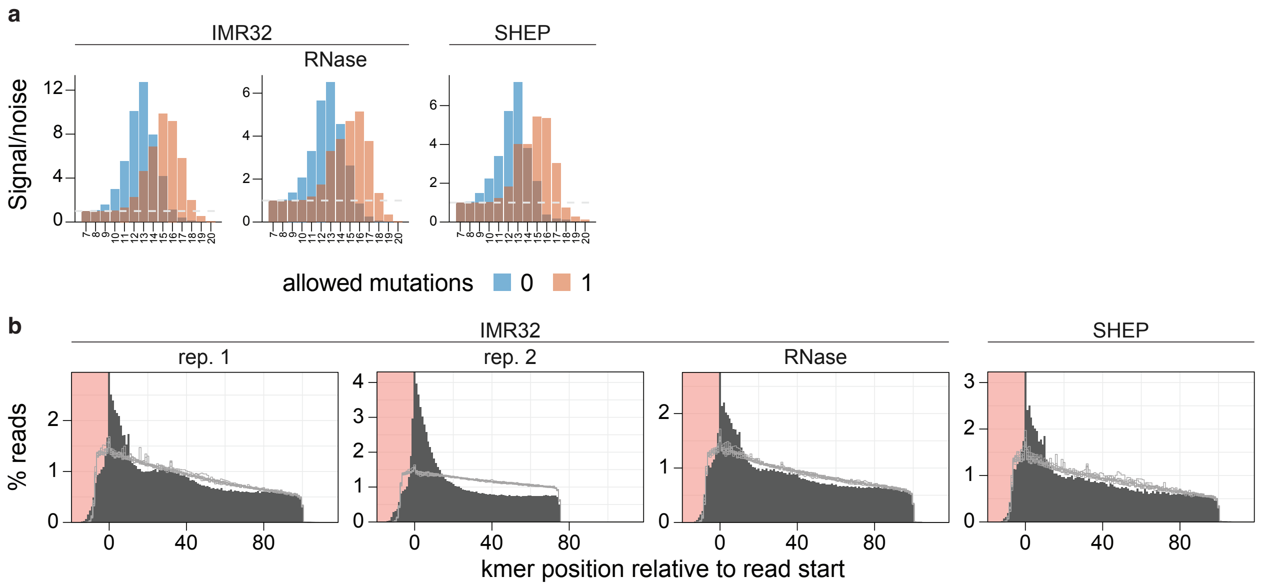


**Figure S8. a,** Signal/noise for the presence of kmers of indicated length in *NESPR* ChIRP-seq peaks from the indicated sample. **b,** enrichment of the longest kmers near ends of reads from the indicated samples in *NESPR* ChIRP-seq data.
